# Polymer functionalized liposomes as universal nanocarriers for drug delivery: Single particle insights on size-dependent performance and intracellular behavior

**DOI:** 10.1101/2025.07.10.664144

**Authors:** Errika Voutyritsa, Athanasios Oikonomou, Georgios Bolis, Vasilis Papageorgiou, Aikaterini Stavroula Vougiatzi, Nikos S. Hatzakis

## Abstract

Nanomedicine requires smart delivery systems that are precise, robust, and universal. While liposomes are established vehicles in drug delivery, their full potential is challenged by limited stability, leakage, insufficient response and limited insight into particle size dependent performance. Here, we provide polymer-modified liposomes (PMLs), engineered for high structural integrity, broad cargo compatibility, and stimuli-responsive cargo release. We thoroughly characterize PMLs at the single particle level shedding light on key structure-function relationships within polydisperse formulations revealing that small vesicles (<100 nm) displayed significantly higher cargo packing densities, while release was independent of vesicle size. PMLs display high versatility effectively encapsulating cargo types ranging from positively or negatively charged small molecules, to oligonucleotides, and proteins. Studies on PMLs interaction with cell membrane show that PMLs maintain high internalization rate in HeLa and hCMEC/D3 brain cells and achieve ∼50% reduction in cell viability within 24 hours when loaded with the anticancer drug 5-fluorouracil. Finaly, PMLs successfully deliver siRNA targeting eGFP in HEK293-d2eGFP cells, achieving a 10-12% knockdown of eGFP expression, as resolved by machine learning-driven single-cell analysis. This work establishes a framework for PMLs high-resolution functional profiling and opens the way for the next generation rational design of tunable PMLs for drug delivery.

## INTRODUCTION

Liposomes stand as leading examples in the design of nanocarriers, offering high versatility in encapsulating both hydrophilic^1-3^ and hydrophobic^4-6^ cargos ranging from small molecules to large proteins.^7^ Yet despite decades of innovation, conventional liposomes continue to face significant limitations such as membrane instability, spontaneous cargo leakage, and minimal responsiveness to environmental cues.^8-11^ Lipid nanoparticles (LNPs) have recently reshaped the field especially in the context of oligonucleotide therapies. Incorporating pH responsive components not only encapsulate nucleic acids but also facilitate the pH response materialising their endosomal escape, setting the stage for the rapid development of the COVID-19 vaccines.^12-14^ However, LNPs remain largely tailored for extremely negatively charged cargos delivery being limited to load oligonucleotide with cytosolic release efficiency rarely exceeding 1-2%. ^8,15,16^

Hybrid architectures -polymer modified liposomes-have been proposed as alternative delivery systems. By integrating stimuli responsive polymers into lipid membranes^17-21^ or forming polymeric cores,^22,23^ PMLs gain structural stability, and response to external triggers e.g. pH or temperature fluctuations releasing their cargo. These delivery systems have been primarily studies in *in vitro* studies and key questions on how size affects the vesicle’s loading across diverse cargo types and release efficiency, or how the polymer modification modulates cellular uptake, and intracellular performance remain unresolved.

Here, we integrated pH responsive polymeric chains into liposome lipid layer and created smart, robust and versatile PMLs intracellular stimuli triggered release of multiple diverse types of biologicals (Fig **1**).

**Figure 1.**
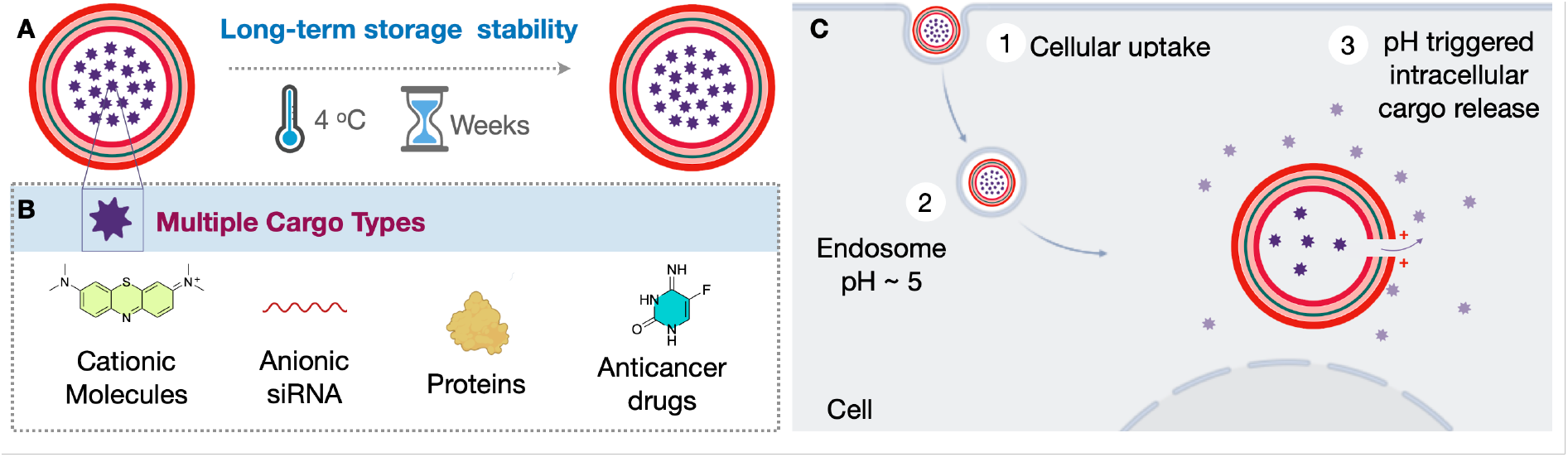
Outline of polymer modified liposomes (PMLs) as a robust, versatile and stimuli-responsive delivery platform. **A**. Cartoon representation of PMLs, loaded with a generic cargo, presenting long term stability as membrane polymer stabilization restrains cargo leakage. **B**. Versatility of PMLs to encapsulate diverse cargo types including cationic, anionic cargos, proteins, and drugs. **C**. Cartoon representation of the PMLs internalization to cells and the pH trigger cargo release due to the pH responsiveness of the incorporated polymer at endosomal acidic pH values. Created with BioRender.com

Combining single particle fluorescence microscopy with quantitative analysis we revealed a hitherto unknown dimensions-to-performance relationships, within PMLs formulation, across polymer and cargo type spanning loading capacity, release efficiency and cellular interaction. We evaluated their delivery efficiency by firstly utilising them to deliver anticancer agents. 5-Fluorouracil (5-FU)-loaded PMLs achieved a ∼50% reduction in viability in HeLa and hCMEC/D3 cells within 24 hours.^24^ GFP-targeting siRNA-loaded PMLs effectively silenced eGFP expression in HEK293-d2eGFP cells. Together, these results establish PMLs as a versatile, efficient and tightly controlled drug delivery platform, capable of precise, cargo-specific delivery across broad medical applications.

## RESULTS AND DISCUSSION

### Robustness of poly(DPA) modified liposomes

Stability and robustness is a critical aspect of any drug delivery system, minimizing uncontrolled and premature drug leakage that results in adverse effects on healthy cells.^25^ To evaluate the impact of polymer modification on the liposome membrane robustness,^18,26^ We decorated both 5% poly(DPA) modified and the non-modified liposome with ATTO 655 on their membrane and loaded them with ATTO 488 carboxy. Membrane integrity and cargo retention over time were tracked on plate reader by measuring ATTO 655 fluorescence (λ_em_= 647 nm) and ATTO 488 carboxy absorbance at 500 nm which was afterwards converted to loaded cargo concentration (Tables S1 and S2). The measurements were taken immediately after the sample preparation and over a 15-day period stored at 4 °C. All samples were dialyzed prior to measurement to ensure that leaked cargo and unbound membrane dye are removed.

Our plate reader assay revealed that 5% poly(DPA) modified liposome retaining almost 95% of the membrane signal and 80% of the loaded cargo after 15 days of storage, whereas the non-modified liposome retained almost 51% and 33% of membrane and cargo signal respectively (Fig. **2A,B**). Polymer modification stabilizes lipid membranes, enabling the design of vesicles with extended shelf-life and superior operational efficiency - key parameters for biomedical applications.

**Figure 2.**
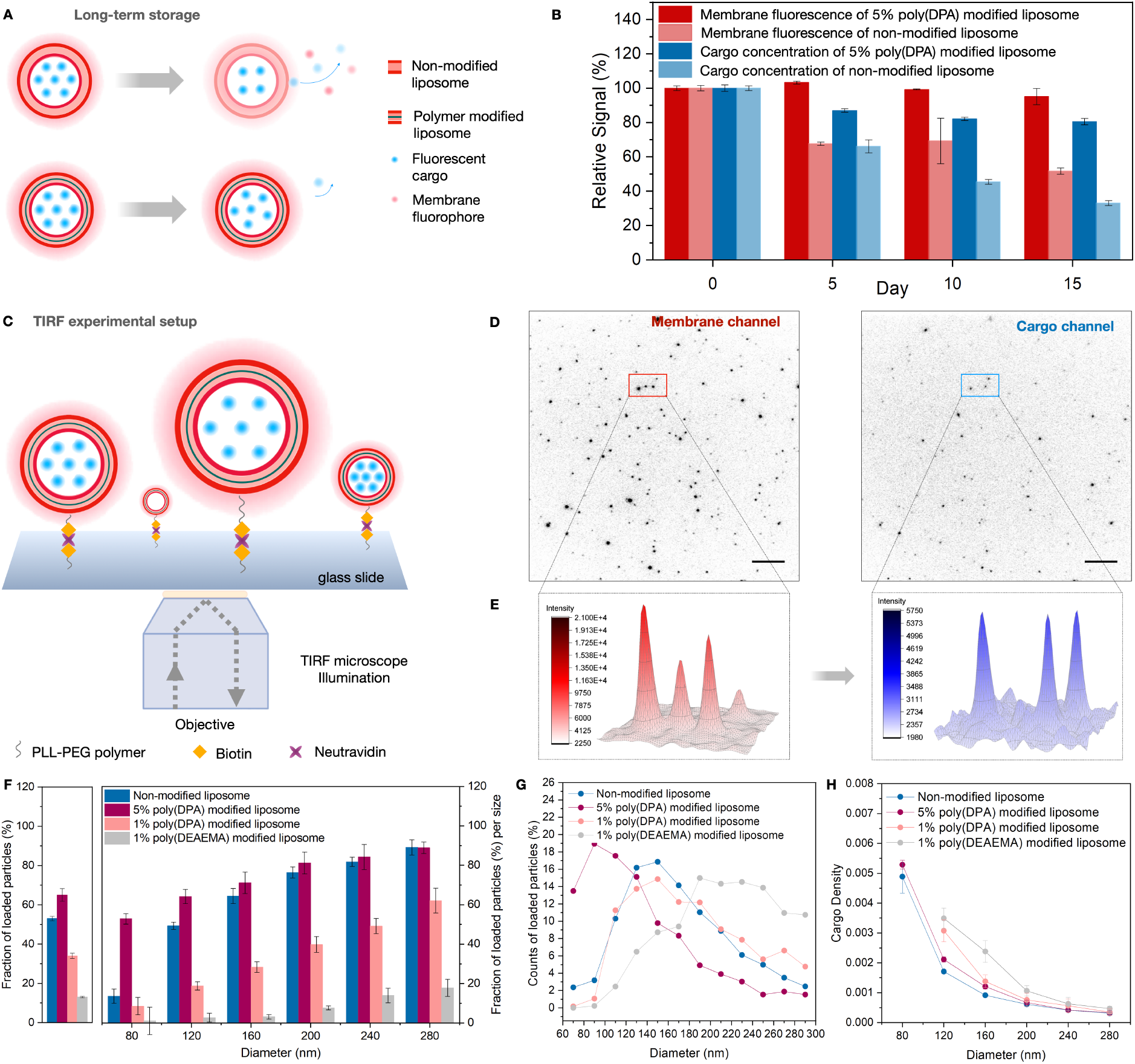
Quantitative comparison of long-term storage stability and encapsulation efficiency of polymer modified and non-modified liposomes. **A**. Cartoon representation of non-modified and 5% poly(DPA) modified liposome labelled with ATTO 655 fluorophore and loaded with ATTO 488 fluorophore losing or maintaining the membrane structure and loaded cargo over time respectively. **B**. Relative membrane fluorescence and cargo concentration of non-modified and 5% poly(DPA) modified liposome at day 5, 10 and 15 as compared to the preparation day stored under dark conditions at 4 °C. Error bars correspond to the standard deviation of three technical replicates of the measurements. The liposome samples were purified with dialysis against 10 mM PBS pH 7.6 before every measurement using dialysis bags MWCO 8-10 kDa. **C**. Cartoon representation of experimental setup on TIRF microscope. Polymer modified liposomes labelled with ATTO 655 fluorophore and loaded with ATTO 488 fluorophore, were tethered on PLL-PEG-passivated surfaces by a biotin-neutravidin linkage. The single particle setup allows observation and quantification of liposome encapsulation variation dependent on particle diameter. **D**. Representative TIRF images of a single field of view imaging thousands of surface-tethered ATTO 488-loaded and ATTO 655-labelled polymer modified liposome. Scale bar corresponds to 1 um. **E**. 3D surface plots of the marked area revealing heterogeneities in size (membrane channel) and encapsulation efficiency (cargo channel) within a population. **F. Left**: Encapsulation efficiency percentage of all the liposome populations. Values are calculated as the ratio of the loaded particles counts to all the detected particles counts of two biological replicates of three technical replicates each. **Right**: Loaded particles percentage of all the liposome populations for each liposome size. Data binned every 40 nm starting from 80 nm. All the liposome populations presented lower percentage of loaded particles at lower sized. Error bars correspond to the standard deviation of the % of empty particles of two biological replicates of three technical replicates each. **G**. Count percentage of loaded particles per bin size for all the liposome population. Data binned every 20 nm starting from 70 nm. **H**. Cargo density of all the liposome populations per bin size. Data binned every 40 nm starting from 80 nm. The data points of 1% poly(DPA) and 1% poly(DEAEMA) modified liposome at 80 nm bin size were excluded representing less that 1% of the particle population of this size range (*see* ***G***). All the liposome populations present a density decrease as the liposome size is increasing, indicating that smaller liposomes are more packed with cargo. Error bars correspond to the standard deviation of the median density of two biological replicates of three technical replicates each. The cartoon representation was created with BioRender.com.

### Single particle studies on loading efficiency of PMLs

To resolve how size heterogeneity shapes PMLs loading efficiency across sample formulations, we capitalised on our single particle assays allowing recordings on thousands of individual PMLs and their cargo synchronously.^27-35^ We produced four PMLs populations starting with non-modified liposomes and 5% poly(2-(diisopropylamino)ethyl methacrylate) (poly(DPA)) modified liposomes that showed significant membrane stability. We also altered the polymer percentage to 1% poly(DPA), and the pH responsive polymer replacing poly(DPA) with poly(2-(diethylamino)ethyl methacrylate) (poly(DEAEMA)), a chemically related polymer. All the PMLs were labelled on their membrane using a fluorescently labelled lipid ATTO-655-DOPE, while ATTO 488 carboxy was used as the fluorescent cargo and were tethered to PEG-passivated microscope surfaces *via* neutravidin-biotin linkage (Fig. **2C**). This methodology enables the parallel imaging and real-time monitoring of each vehicles’ membrane (ATTO 655) and cargo (ATTO 488) signal, offering quantification of the dependence on loading and stability on their dimensions (Fig. **2D,E**).^31^

Our findings revealed that among the liposome populations, 5% poly(DPA) modified vesicles showed higher encapsulation efficiency (∼65%) (Fig. **S1**). This can be attributed to the partial membrane positive charge (∼8% protonation of poly(DPA) amine groups at pH 7.6) promoting electrostatic interactions with the weekly anionic cargo. Reducing the polymer content to 1% diminished the effect leading to lower loading capacity. In contrast, despite its higher membrane positive charge (∼20%), poly(DEAEMA) modified liposomes showed poor encapsulation efficiency likely attributed to reduced membrane stability at physiological pH as the pKa of poly(DEAEMA) is close to 7.0 (Fig. **2F**, left).

Although all formulations were extruded through 100 nm pore size membrane, individual vesicle sizes ranged broadly from 50 nm to 300 nm, consistent with our earlier studies,^31,33,35^ revealing substantial sample heterogeneity that remains masked in conventional readout averaging the behavior of a large number of vesicles. Our single particle readout allowed us to directly observe each vesicle as an autonomous particle where the dimensions and the amount of loaded cargo can be quantified. Single-particle vesicle sizes were extracted by mapping background-corrected fluorescence intensities to physical dimensions, considering their spherical geometry. Intensities were extracted and translated into absolute size (nm) via calibration against the average vesicle diameter measured on DLS, matching the square root of the mean integrated intensity to the DLS-derived diameter. This yielded a calibration constant, enabling high-confidence intensity-to-dimension transformation at single-particle resolution.^31^ To correlate the dimension of every single liposome to its capacity to load cargo, we extracted the background-corrected intensities from the colocalized cargo channel for each membrane-defined vesicle. The single particle readout allowed parallel cargo intensities across thousands of individual liposomes, which in turn revealed a clear correlation between liposome size and cargo loading. Using 40 nm size binning, we classified vesicles into size groups. Across all sizes 5% poly(DPA) modified liposomes consistently exhibited the highest fraction of loaded particles, while unloaded vesicles were systematically measured smaller, clustering around 50 nm across all populations (Fig. **2F**, right and Fig.**S2**).

To reveal how polymer modification and density reshape size-dependent loading profile, we performed single-particle analysis decreasing the binning size to 20 nm to achieve higher-resolution mapping with smoother, well-fitted distribution curves. Smaller sizes of 5% poly(DPA) modified liposomes presented higher loaded vesicle percentage (peaked at ∼90 nm) compared the non-modified and 1% poly(DPA) modified liposomes (peaked at ∼150 nm). In the case of 1% poly(DEAEMA) the peak was shifted to ∼190 nm, highlighting how polymer structure and density reshape size-dependent loading behaviour (Fig. **2G**). Cargo intensity scaled with size, as expected, but when normalized to vesicle volume, smaller vesicles demonstrate significantly higher cargo density (Fig. **2H** and Fig. **S3**), indicating more efficient packing. Our findings suggest that vesicle size may influence not only loading but also the cargo packing within vesicles, a feature crucial in therapeutic applications requiring compact delivery.

### Stimuli triggered cargo release rates of PMLs by single particle studies

To assess the pH response of PMLs under endosomal-like environment, we directly monitored the membrane and cargo signal upon acidification from 7.6 to 5.2, using our dual-color single particle assay and imaging every 30 minutes over a two-hour period (Fig. **3A,B**). Our methodology enabled the parallel imaging across thousands of individual PMLs tethered on the microscope surface.

**Figure 3.**
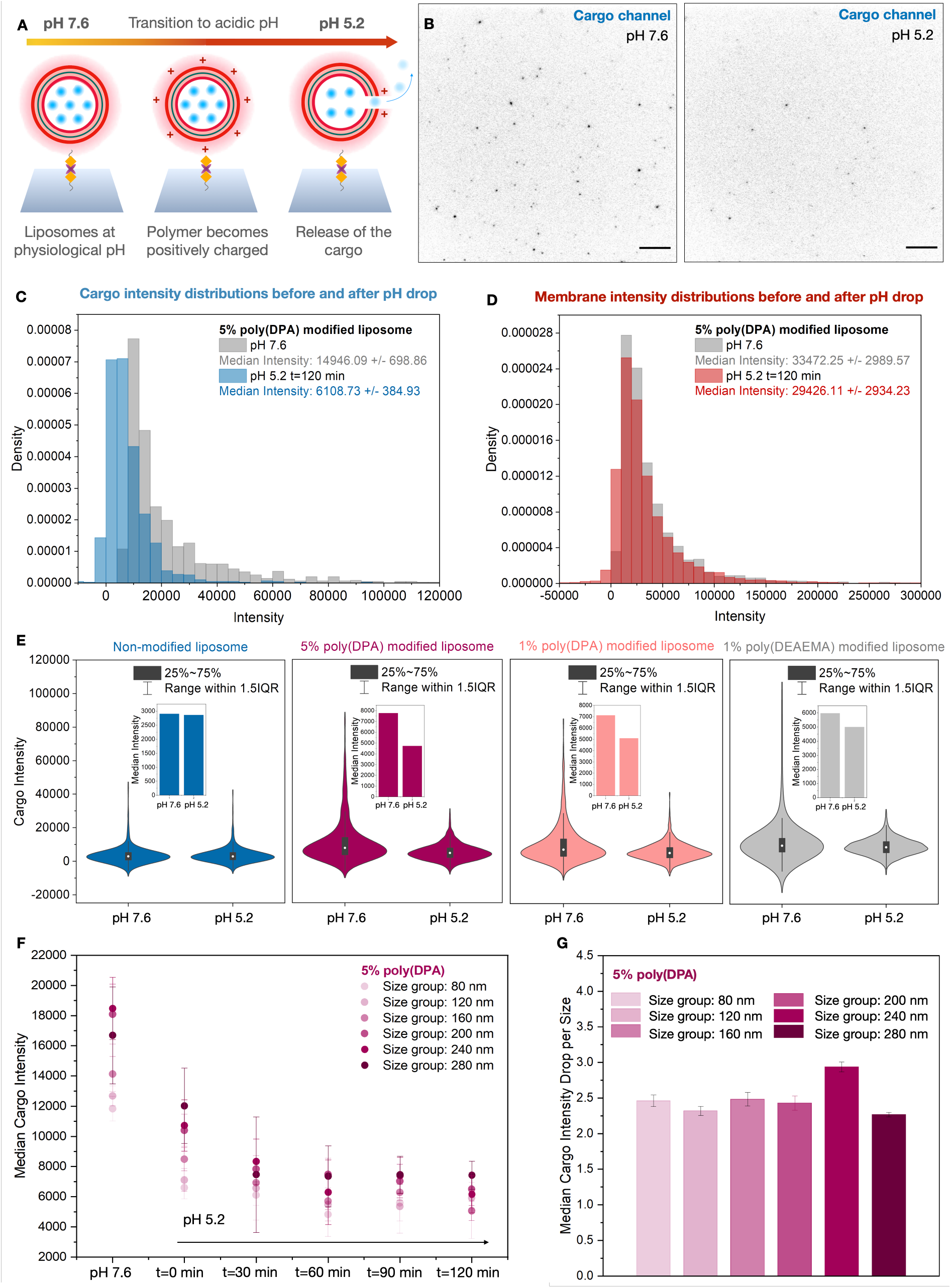
Direct observation and quantification of extent of stimuli triggered cargo release of polymer modified liposomes *in vitro*. **A**. Cartoon representation of experimental setup on TIRF microscope. Polymer modified liposomes labelled with ATTO 655 fluorophore and loaded with ATTO 488 fluorophore, were tethered on PLL-PEG-passivated surfaces by a biotin-neutravidin linkage. The single particle setup allows observation and quantification of the cargo release upon pH drop from physiological to acidic pH. **B**. Representative TIRF images of a single field of view imaging thousands of surface-tethered ATTO 488-loaded polymer modified liposome at pH 7.6 and pH 5.2 incubated for 120 min, displaying signal loss due to cargo release. Scale bar corresponds to 1 μm. **C**. Background corrected intensity distributions of 5% poly(DPA) modified liposome on the cargo channel at pH 7.6 and at pH 5.2 after 120 min incubation at acidic pH. A drop in pH from 7.6 to 5.2 results in reduction of the median cargo intensity distribution by 59%. Error bars correspond to the standard deviation of two biological replicates. **D**. Background corrected intensity distributions of the membrane channel of 5% poly(DPA) modified liposome shown negligible intensity decay from pH 7.6 to pH 5.2 after 120 min incubation (from 33472 to 29426), supporting that cargo loss does not originate from loss of liposomes structural integrity. **E**. The cargo intensity range of non-modified (blue), 5% poly(DPA) modified (magenta), 1% poly(DPA) modified (pink) and 1% poly(DEAEMA) modified (grey) liposome at pH 7.6 and 5.2. Bar charts insert: Median Intensity values at pH 7.6 and pH 5.2 for each liposome population. **F**. Median cargo intensity values of the 5% poly(DPA) modified liposome at pH 7.6 and at pH 5.2 at 5 time points from 0 to 120 min per size. Data binned every 40 nm starting from 80 nm. Error bars correspond to the standard deviation of the median intensities of two biological replicates. **G**. Median cargo intensity drop from pH 7.6 to pH 5.2 after 2 hours incubation per size. Error bars correspond to the standard deviation of the median intensities of two biological replicates. The cartoon representation was created with BioRender.com.

All PMLs formulations displayed a pronounced cargo signal loss immediately after acidification, followed by a gradual decrease over two hours (Fig. **3C**, and Fig. **S4**-**S7**). In contrast, membrane signal remained stable, confirming that the observed cargo loss did not result from vesicle rupture but from membrane destabilization (Fig. **3D**, and Fig. **S10, S11**). Quantitative analysis of the entire vesicle populations revealed that 5% poly(DPA) modified liposomes exhibited the highest overall cargo signal reduction (∼40% total loss), followed by 1% poly(DPA) (∼29%) and 1% poly(DEAEMA) (∼16%) (Fig. **3E**). Control experiments verified that photobleaching was not accounted for the intensity loss in the cargo channel. In comparison, focusing on loaded vesicles only, cargo signal loss reached ∼60% for 5% poly(DPA), ∼62% for 1% poly(DPA) and ∼77% for 1% poly(DEAEMA) liposome population suggesting that polymer, rather than density, more strongly affects per particle release efficiency. The slightly higher cargo release for poly(DEAEMA) per loaded particle can be attributed to its higher pKa (7.0), rendering the membrane more susceptible to protonation and destabilization at acidic pH compared to poly(DPA) (6.5). As expected, non-modified liposome presented minimal cargo signal loss (1-3%), highlighting the polymer-driven stimuli responsiveness of the PMLs.

Single particle mapping and quantitative analysis of the recorded membrane and cargo readouts revealed that all particle sizes respond to pH drop and release their cargo (Fig. **3F** and Fig. **S8**). The decay rates of each size group were also identified and revealed similar release efficiency for all the particles regardless of their size (Fig. **3G**).

### PMLs as a universal nanocarrier

To assess the versatility of PMLs, we evaluated the encapsulation of chemically diverse cargos in 5% poly(DPA) modified liposomes: methylene blue (slightly cationic), siRNA (highly anionic), and BSA (large protein) (Fig. **4A**). Despite the vesicle membrane partial positive charge at physiological pH, the methylene blue achieved the highest loading fraction (∼70%), indicating that electrostatic repulsion it’s not an encapsulation limitation factor (Fig. **4B**). The highly negatively charged siRNA reached ∼55% loading, while the loading of a protein was also successful with bovine serum albumin (BSA) to show ∼40% loading efficiency, likely limited by its large molecular size (Fig. **4B**). Single particle analysis revealed that methylene blue and siRNA loaded vesicles are dominant across all size bins, while BSA loaded vesicles showed significantly lower probability in smaller sizes (<160) likely due to steric hindrance (Fig. **4C**), suggesting that both cargo charge and size affect PMLs loading efficiency.

**Figure 4.**
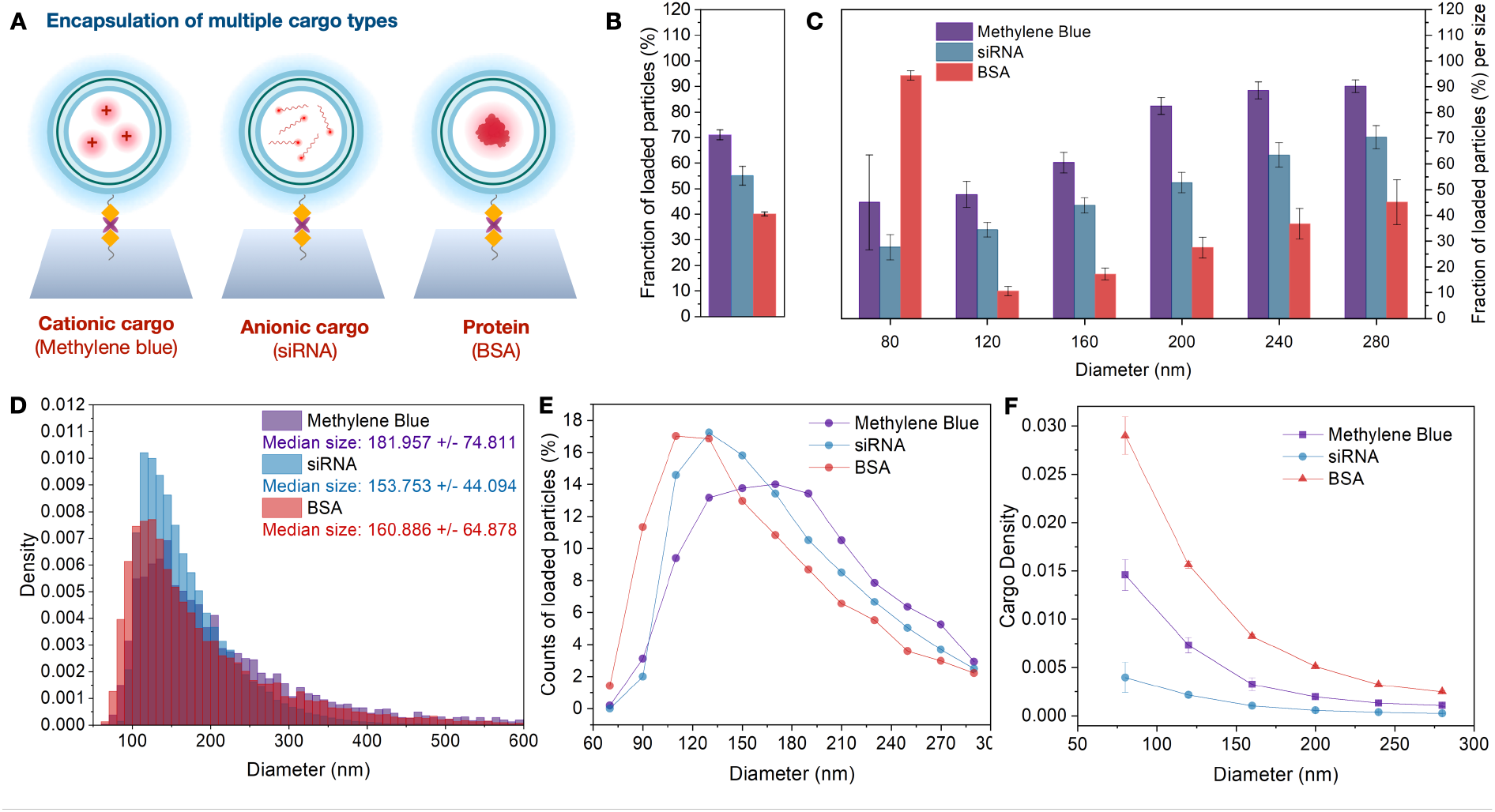
Polymer modified liposome as a versatile platform to efficiently encapsulate diverse cargo types. **A**. Cartoon representation of polymer modified liposomes loaded with 3 types of cargo: Methylene Blue as a cationic dye, siRNA as a multiple anionic nucleic acid, and Bovine Serum Albumin (BSA) as a protein, tethered on PLL-PEG-passivated surfaces by a biotin-neutravidin linkage. **B**. Encapsulation efficiency percentage of the 5% poly(DPA) modified liposome loaded with the three different cargo types. Values are calculated as the ratio of the loaded particles counts to all the detected particles counts of six technical replicates. **C**. Loaded particles percentage of the 5% poly(DPA) modified liposome loaded with the three different cargo types for each liposome size. Data binned every 40 nm starting from 80 nm. All the liposome populations presented lower percentage of loaded particles at lower sized. Error bars correspond to the standard deviation of the % of empty particles of six technical replicates. **D**. Size distribution of the 5% poly(DPA) modified liposomes loaded with the three different cargo types, indicating that the cargo type only marginally affects the size of the formatted particles. **E**. Count percentage of loaded liposomes per bin size for methylene, blue, siRNA, and BSA loaded 5% poly(DPA) modified liposome. Data binned every 20 nm starting from 70 nm. For methylene blue most of the loaded liposomes are detected at the 170 nm bin size, for siRNA at the 130 nm bin size and for BSA at 110 nm bin size, indicating that the loading process is affected from the cargo type. F. Cargo density of the 5% poly(DPA) modified liposome loaded with the three different cargo types at each bin size, indicating that smaller liposomes are more packed with cargo in all cases. Error bars correspond to the standard deviation of the median density of six technical replicates. The cartoon representation was created with BioRender.com.

Size distribution confirmed similar dispersion across the three populations, with median sized between 150-180 nm (Fig. **4D**). However, differences in the loaded particle counts per size bin peak were pronounced, as the repulsive electrostatic forces present in methylene blue loaded vesicles sifted the peak to larger sizes of 150-190 nm, while vesicles loaded with siRNA and BSA peaked at 130 and 110 nm respectively (Fig. **4E**). Quantification of the cargo signal per particle and normalization to vesicle volume, revealed distinct packing behaviours (Fig. **4F**, and Fig. **S3**). Methylene blue showed low cargo density overall, while siRNA loaded vesicles maintained consistent packing over a narrow size range. BSA loaded liposomes, despite their lower loading efficiency and large cargo size, exhibited the highest cargo density and the higher fraction of loaded particles at smaller sizes.

### Single cell studies on PMLs internalisation rate and intracellular delivery

Drug delivery nanoparticles that incorporate poly(DPA), such as polymerosomes, have shown promise in targeting tumor cells, enhancing cellular uptake and delivery of therapeutic agents e.g. doxorubicin.^37^ At the same time, liposomes modified with pH-sensitive polymers exhibit improved cellular uptake and controlled release of encapsulated drugs when delivered in cancerous tissues.^38^ Here, we extend these findings by demonstrating that poly(DPA) modified liposomes efficiently internalize and release their cargo inside epithelial and endothelial cells.

To assess the cellular delivery potential of 5% poly(DPA) modified liposomes, we conducted single particle tracking (STP) studies^31,32,34,39-43^ and captured their interaction with cells. Image analysis through our custom single particle tracking toolboxes allowed us to directly observe the spatiotemporal localization of thousands of individual PMLs interacting with HeLa and hCMEC/D3 cells.^32,44^ Dual-colour parallel recordings allowed tracking in the red channel liposomes labelled with ATTO 655 on their membrane and HeLa and hCMEC cells in the blue channel with ATTO 488 as the dye exclusion marking the external solution around the cells (Fig. **5A**). Cell boundary identifying was accomplished using our custom analysis algorithms Our results showed a time-dependent increase in PMLs interaction with cells for both cell lines (Fig. **5B**) indicating that the polymer modifications does not prevent the interaction with cell membrane. In summary these findings demonstrate the efficiency of PMLs in the cellular uptake process, and their potential for drug delivery applications in targeting hard-to-transfer cell types.

**Figure 5.**
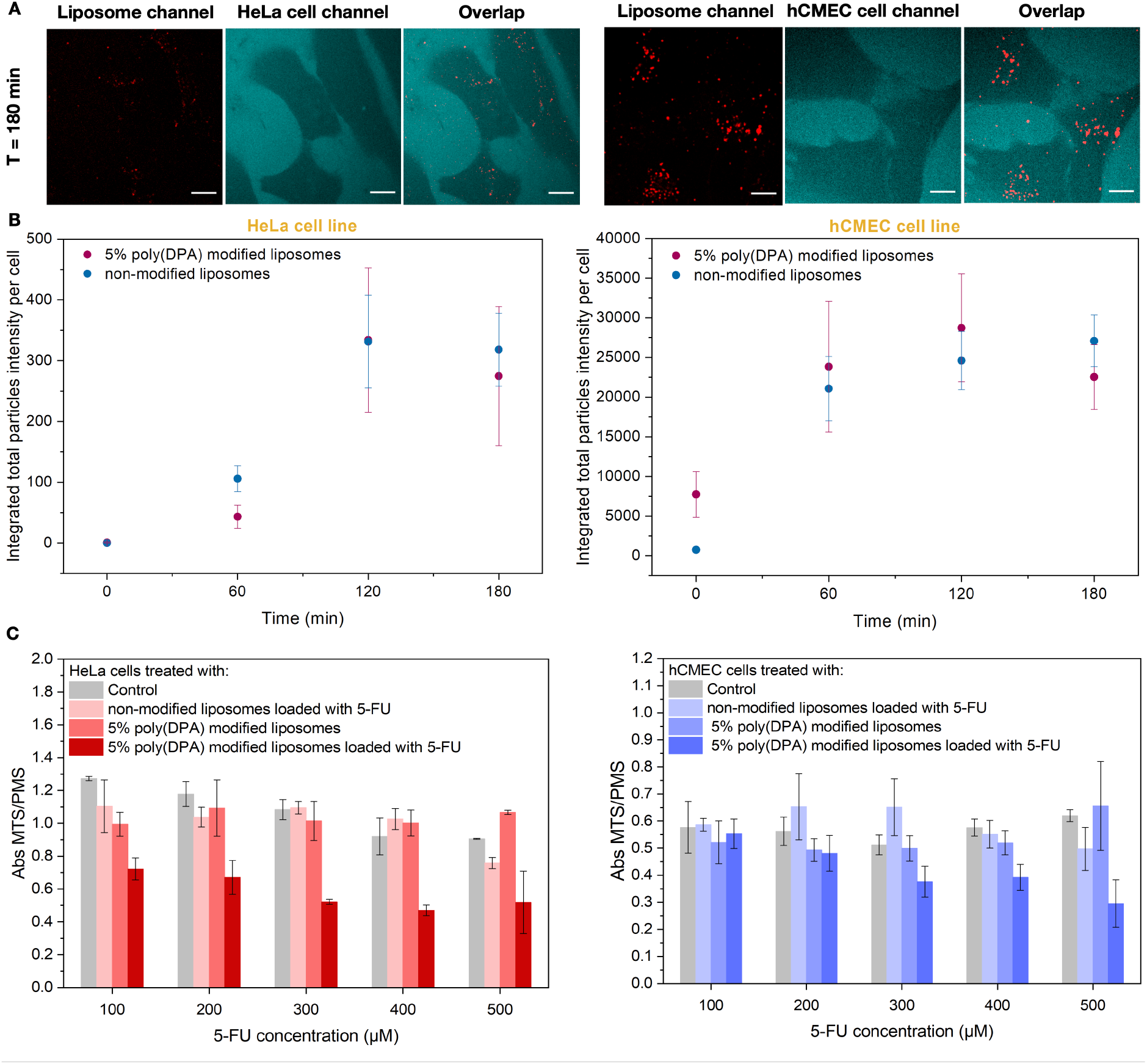
Direct observation of spatiotemporal localization of liposome internalized in HeLa cells and hCMEC/D3 endothelial cells. **A**. Representative 2D images acquired by spinning disc microscopy showing 5% poly(DPA) modified liposomes being either stuck on the cell membrane or internalized after 3 hours incubation. Addition of ATTO 488 carboxy dye offers identification of HeLa and hCMEC/D3 cells that appear darker, liposomes are labelled with ATTO 655. Scale bar is 20 μm. **B**. Total particles’ intensity (membrane bound or internalized) per cell over time for HeLa cancer and hCMEC/D3 brain cell lines treated with either the non-modified or 5% poly(DPA) modified liposome. Error bars correspond to the standard deviation of the mean cell intensity for 10 fields of view. The mean number of analysed cells was 18 for HeLa cells treated with 5% poly(DPA) modified liposomes, 10 HeLa cells treated with non-modified liposomes, 9 hCMEC/D3 cells treated with 5% poly(DPA) modified liposomes, and 8 hCMEC/D3 cells treated with non-modified liposomes. Cell segmentation was performed with Cellpose software.^36^ **C**. MTS-PMS assay for cell viability for different concentrations of 5-FU loaded in 5% poly(DPA) modified and non-modified liposomes. Data are recorded 24h after liposome incubation in HeLa and hCMEC/D3 cells. Error bars correspond to the standard deviation of three technical replicates of the measurements.

To evaluate the intracellular release and the potential of PMLs as efficient drug delivery nanocarriers we measured the cytotoxicity of 5% poly(DPA)-modified liposomes loaded with the anticancer drug 5-fluorouracil (5-FU) applying standard MTS-PMS assay in HeLa and hCMEC/D3 cells. The method relies on the reduction of tetrazolium salt (MTS) to a formazan dye by viable cells in the presence of an electron-coupling reagent (PMS). The results demonstrated a dose-dependent cytotoxic response, indicating effective delivery of 5-FU. In HeLa cells a 300 μM 5-FU dose resulted in cell viability by approximately 50% after 24-hour incubation. A higher dose of 500 μM required for hCMEC cells also exhibiting susceptibility to the cytotoxic anticancer drug (Fig. **5C**). Besides the efficient internalization PMLs display functional drug release within two types of cells, demonstrating their potential for drug delivery application.

To further assess the universality of PMLs, we evaluated their capacity to deliver oligonucleotides, widely considered as next generation pharmaceutics^12,13^ and the central scope of multiple research groups.^45-48^ An established method to evaluate efficient siRNA delivery in cells is the eGFP protein expression knockdown resulting in signal loss due to degradation of mRNA translated to the fluorescent protein, upon siRNA delivery.^49,50^ For microscopy applications, the destabilized variant of eGFP (d2eGFP) is widely used due to the fast turnover (2 hours half-life) enabling direct observation of the fluorescent signal attenuation after treatment with siRNA.^51^ Here, we assessed the intracellular functionality and eGFP targeting efficiency of siRNA loaded PMLs in HEK293-d2eGFP cells, by imaging HEK293-d2eGFP cells on spinning disk microscope under two conditions: untreated cells as the control experiment and cells treated with 5% poly(DPA) modified liposomes loaded with eGFP-targeting siRNA (Fig. **6A**). Imaging the cell volume with a frame rate of 2.5 min for 10 h ensured minimal photobleaching and photodamage with prolonged imaging to observed directly eGFP signal decay.

**Figure 6.**
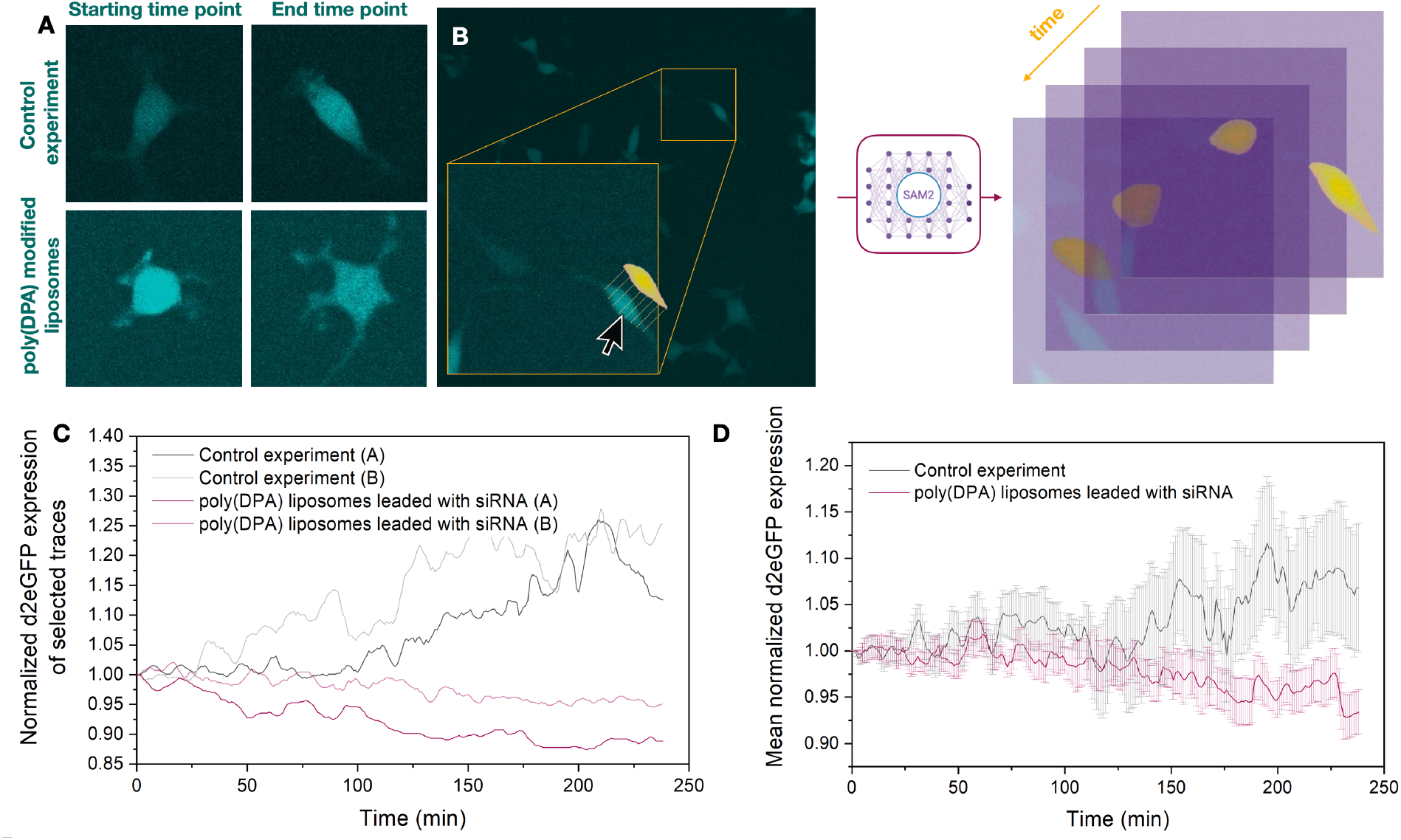
Direct observation of d2eGFP knockdown in HEK293 cells treated with siRNA targeting d2eGFP loaded PMLs. **A**. Representative 2D images acquired by spinning disc microscopy showing that start and end point of HEK293 cells expressing d2eGFP in the absence of liposomes (control experiment) and treated with 5% poly(DPA) modified liposomes loaded with siRNA targeting d2eGFP over a ten-hours period of time. HEK293 cells are imaged using a 488 nm laser for the excitation of the d2eGFP. **B**. Single cell analysis for d2eGFP level quantification pipeline: A cell of interest is selected and segmented using Cellpose.^36^ The cell mask is used as prompt for SAM2 (Segment Anything Model 2)^52^ and the cell is tracked over time. **C**. Two representative traces normalised for each of the above experimental condition. In the absence of particles HEK293 cells increasingly express d2eGFP over time, while PMLs delivered siRNA results in 10% knockdown. Each time point intensity represents the median intensity of the cell mask normalized to the mean value of the 3 first points of a trace. Intensity traces were denoised with a uniform convolution window of length 5. **D**. Median intensity of d2eGFP expressed HEK293 cells of all the selected cells for each experimental condition over time. The errors are the standard deviation of the mean intensity of the selected cells.

To quantify the dynamic changes in eGFP intensity, we employed Segment Anything Model 2 (SAM2),^52^ a transformer-based, prompt-driven model to segment cells. Cells showing minimal overlap with neighbouring cells were selected and cell masks were created at the starting time point. These masks were tracked across timepoints, enabling the extraction of fluorescence intensities per selected cell across time, generating time dependent trajectories of eGFP signal of individual cells (Fig. **6B**). Our analysis revealed that in the control experiment cells exhibited increasing eGFP signal with a mean rate of +3.42 × 10^−4^ A.U. per minute, while 5% ply(DPA) modified liposomes loaded with eGFP-targeting siRNA effectively delivered their cargo in HEK293 cells resulting in 10-12% d2eGFP rate knockdown with a mean rate of -2.46 × 10^−4^ A.U. per minute, confirming functional siRNA delivery (Fig. **6C,D** and Fig. **S15**). At the single cell level, 50% of the selected tracked cells in the control experiments showed increasing eGFP signal over time, while upon treatment with 5% ply(DPA) modified liposomes loaded with eGFP-targeting siRNA, 75% of the selected tracked cells showed a clear eGFP signal decrease. Our results demonstrate the intracellular siRNA delivery and gene silencing outcomes of responsive PMLs achieving more than 10% mean d2eGFP knockdown.

## CONCLUSIONS

This study establishes PMLs as a robust, versatile, and stimuli-responsive nanocarrier platform for precise drug delivery. By combining advanced fluorescence microscopy with custom particle tracking algorithms, we evaluated how their dimensions, polymer density, and cargo type alter nanocarrier performance resolving structure-function relationships that are masked in conventional assays. Poly(DPA)-modified liposomes exhibited enhanced membrane stability, reduced uncontrolled leakage, and efficient pH-triggered release while maintaining enhanced structural integrity as compared to the conventional liposome counterparts. Smaller vesicles displayed increased packing density, though cargo release rates were size-independent, indicating consistent cargo release behavior across population. Importantly, PMLs demonstrated effective intracellular delivery, as confirmed by their high interaction with both epithelial and endothelial cancer cell membranes, and by the successful cytosolic delivery of nucleic acid cargo achieving siRNA-mediated gene silencing and ∼50% reduction in cell viability with 5-FU payloads.

These findings introduce PMLs as efficient delivery vehicles, capable of diverse therapeutic cargos into cells. Besides showcasing the value of single-particle functional mapping in rational nanocarrier design, this study paves the way for next-generation, stimuli-responsive delivery systems tailored for intracellular targeting and personalized therapy.

## EXPERIMENTAL METHODS

### Functionalized liposome preparation

The desired mole percentage of lipids was mixed in a glass vial. The glass vial was dried by softly flowing pure nitrogen for 10 -15 minutes until the lipids formed a thin film. The vial was furthermore vacuumed for at least one hour. The lipids were rehydrated to a lipid concentration of 1g/L with 10 mM PBS buffer pH 7.6 and vortexed for 30 seconds. The liposome sample were incubated for 60 minutes to self-assemble. The liposome sample were flash-frozen in a mixture of dry ice and acetone followed by thawing in a heat bath. This was repeated 10 times to ensure unilamellarity. To ensure homogeneous size distribution, the liposome sample were extruded 10 times (back and forth) through a 100 nm pore size membrane. The liposome sample were aliquoted and stored at -20 ^°^C (*see Supplementary Information for the lipid composition of each functionalized liposome population*).

### Substrate surface preparation for single particle imaging

Glass slides 76 × 26 nm were dried with a nitrogen flow and activated using a plasma cleaner for 1 minute at a pressure between 300 mtorr to 400 mtorr. Immediately after activation the slides were attached to the glass slides to make flow cells. Each of the six champers was passivated with 80 μL of a PLL-g-PEG (1 mg/mL in HEPES buffer pH 5.5) and PLL-g-PEG-biotin (1 mg/mL in HEPES buffer pH 5.5) mixture in the ratio 100:1. The surface was incubated for 30 min. Excess of PLL-g-PEG mixture was removed by flushing five times with 50 μL HEPES buffer pH 5.5. 80 μL of the neutravidin solution (1 mg/mL in HEPES buffer pH 5.5) was added in each well. The surface was incubated for 15 min. Each well was washed with 10 mM PBS buffer pH 7.6 at least 5 times before the sample addition.

### Single particle assay for loading and release experiments

A TIRF microscope (IX83, Olympus) was used for the single-particle imaging experiments. Oil immersion objective (UAPON 100XOTIRF, NA 1.49, Olympus) and an EMCCD camera (ImagEM X2, Hamamatsu, Shizuoka, Japan) were used to record images and videos with a pixel width of 160 nm and field of view 81.92 μm × 81.92 μm. Laser lines of 640 nm and 488 nm were used to excite the fluorophores ATTO-655-DOPE (liposome membrane) and ATTO-488 carboxy (cargo), respectively. Imaging was performed with an exposure time of 50 ms, 100 nm penetration depth, and 300 EM gain. Each image series with a frame rate 2 frames per 5 seconds, contained 10 frames of the 488 nm channel and 10 frames of the red channel alternating between them. Prior to imaging the sample (C = 1.5 10^−2^ mg/mL) was incubated for 10 minutes and washed with 10 mM PBS buffer pH 7.6 (6 × 0.5 mL) for liposome immobilization. For the release assay, after imaging at pH 7.6, the buffer solution was exchanged to 156 mM sodium acetate buffer pH 5.2. Each well was washed at least 5 times before every measurement with the buffer solution of the chamber for the fluorophore that is released in the solution to be removed.

### Calculation of membrane integrity and ATTO-488 carboxy leakage over time

5% poly(DPA) modified and non-modified liposome populations labelled with ATTO 655 and loaded with ATTO 488 carboxy (10^−2^ μmol/mL) in 10 mM PBS pH 7.6 were prepared. The fluorescence of ATTO 655 at 647 nm and the absorbance of ATTO-488 carboxy at 500 nm was measured immediately after preparation and after 5, 10 and 15 days of storage at 4 °C. The absorbance measurements were converted to ATTO-488 carboxy concentration. The liposome populations were purified via dialysis in 10 mM PBS pH 7.6 before every measurement for the leaked ATTO-488 carboxy to be removed.

### Cell culture

HeLa and HEK293 (stably expressing d2eGFP) cells were cultured in Nunc EasYFlask 25 cm^2^ cell culture flasks with Dulbecco’s Modified Eagle’s Medium with 4500 mg/L glucose supplemented with 2 mM L-glutamine, 1 mM sodium pyruvate, and 10% (v/v) heat-inactivated FBS. hCMEC/D3 cells were cultured in EBM-2 endothelial basal medium (Lonza) supplemented with 2mM L-glutamine and 10% (v/v) heat-inactivated FBS. Both cell lines were detached by 3 × trypsin-EDTA treatment (5 mg/mL trypsin and 2 mg/mL EDTA in DPBS, pH 7.4) and seeded for subsequent experiments. Cells were incubated in a humidified, 5% CO_2_, 37°C incubator. The passages for all cell experiments ranged from 7 to 14.

### Single cell assay for liposome internalization

An oil immersion 60 x objective (Olympus) and a numerical aperture of 1.4 connected to a CMOS camera (photometric PRIME 95B) with an effective pixel size of 183 nm × 183 nm was used for cell imaging. Cells were grown into 8-well plates 24h before imaging. After incubation, the culture media was removed, and the cells were washed twice with PBS. Subsequently, 200 μL of imaging media and 0.4 μL of ATTO 488 Carboxyl 50 mM dye exclusion were added to each well, followed by the addition of 1 μL of liposome (membrane-labelled with ATTO 655) solution of a concentration of 1 mg/mL. For each experimental condition, 10 positions were imaged in parallel directly after the addition of the liposomes and every 1h in a 3h-time period. Imaging was performed with an exposure time of 50.04 ms. Each image series with a frame rate 2 frames per 1 seconds contained 100 frames of the 488 nm channel and then 100 frames of the 647 nm channel. In the acquired images series, cells were tracked over time using an in-house developed tracking algorithm.

### Single cell assay for d2eGFP knockdown

HEK293 cells stably expressing d2eGFP were established by transducing cells with a lentiviral vector encoding d2eGFP. The lentiviral particles were produced by transfecting the HEK293T packaging cell line with pMD2.G, psPAX2, and pLV-CMV-d2eGFP plasmid using Lipofectamine 3000 according to the manufacturer’s protocol. HEK293 cells stably expressing d2eGFP were transferred to a preheated microscopy incubation chamber, and 5 positions with evenly distributed cells were selected for each experimental condition (cells in the absence of particles, and cells treated with siRNA loaded 5% poly(DPA) modified liposomes). Immediately before starting image acquisition, of 1 μL of liposome solution of a concentration of 1 mg/mL loaded with siRNA targeting d2eGFP were added dropwise to the medium of the wells of interest. Seven z-plane images with 3 μm z-spacing were acquired per position at 2.5 min intervals. Images were acquired for 10 h in total. For the excitation of the d2eGFP, 488nm laser was used with a 50,04 ms exposure time. **MTS-PMS assay:** For the cytotoxicity studies 4 liposome populations were prepared: 5% poly(DPA) modified liposome population loaded with the anticancer drug 5-FU (3mM), non-loaded 5%poly(DPA) modified liposomes and non-modified liposomes loaded with5-FU (3mM) in 10 mM PBS pH 7.6. HeLa and hCMEC/D3 cells were seeded in a 96-well plate at a density of 1.5 × 104 cells per well and were incubated in Dulbecco’s Modified Eagle’s Medium for 24 h at 37 °C. Liposomes loaded with 5-FU were dialyzed against 10 mM PBS pH 7.6 for the unloaded 5-FU to be removed. All the liposome populations were added to the medium and the cells were incubated at 37 °C for 48 h. The drug concentrations were set between the range 100-500 μM. The nanoparticle concentrations were also varied according to the drug concentrations. Cell viability was determined with the CellTiter 96® AQueous Non-Radioactive Cell Proliferation Assay (Promega) following the manufacturer’s protocol.

### Image Analysis

For the intensity extractions, an in-house, GPU accelerated, Laplacian of Gaussian was used for particle detection and intensity extraction. Nearest Neighbours was implemented on the GPU accelerated framework for particle tracking. Custom Python scripts were developed and used for the post-detection intensity analysis and statistics for both static and temporal experiments. The radius of the liposomes was determined by formulas demonstrated in Quantifiable Information on a static image subsection. For each particle, the intensity within the ROI (defined by the σ of its detection) was integrated and adjusted for background noise. For GFP transfection experiments on HEK293 cells, cell segmentation and tracking were performed using a foundational mode (SAM2)^52^ with bounding boxes on cells of interest.

## AUTHOR INFORMATION

### Author Contributions

E.V., with inputs from NSH, conceived of the project idea and designed the project.; E.V. produced all the liposome formulations, performed all the stability and characterization measurements in bulk and all the single particle experiments on TIRF microscope; E.V., G.B., and V.P. performed the experiments on cells; A.O. wrote the script for particle and cell intensity extraction; E.V. and A.O. carried out the data analysis; N.S.H. had the overall supervision, project management and provided resources; E.V. wrote the original draft of the manuscript; A.O. edited the manuscript; N.S.H. reviewed, edited and rewrote parts of the manuscript; All authors have read and agreed to the published version of the manuscript.

## ACKNOWLEDGMENT

The authors acknowledge funding from the Independent Research Fund Denmark (1127-00432B), the NNF challenge Center for Optimised Oligo Escape and control of disease (NNF23OC0081287), the NNF Center for 4D Cellular Dynamics (NNF22OC0075851), the Villum foundation Synergy grant (40578), and the Villum experiment grant (40801).

