## Supplementary Information for "Polymer functionalized liposomes as universal nanocarriers for drug delivery: Single particle insights on size-dependent performance and intracellular behavior"

a. Department of Chemistry, University of Copenhagen, Denmark
b. Nanoscience centre, University of Copenhagen, Denmark
c. Novo Nordisk center for Optimized Oligo Escape, University of Copenhagen, Denmark
d. Center for 4D cellular dynamics, Department of Chemistry, University of Copenhagen, Denmark

### 1. Materials

Chemicals such as phosphate buffer saline (product number: P4417), sodium acetate (CAS: 127-09-3), acetic acid glacial (CAS: 64-19-7), 5-fluorouracil (CAS: 51-21-8), cholesterol (CAS: 57-88-5), poly(2-(diisopropylamino)ethyl methacrylate) (poly(DPA), average $M_n$: 10.000, product number: 910457), poly(2-(diethylamino)ethyl methacrylate) (poly(DEAEMA), average $M_n$: 10.000, (product number: 910104), Methylene Blue (CAS: 122965-43-9), ATTO 655 NHS ester (product code: 76245), and Bovine Serum Albumin (BSA) (CAS: 9048-46-8) were purchased from Sigma-Aldrich. Hydro Soy PC (HSPC, CAS: 97281-48-6), DSPE-PEG$_{(2000)}$ Biotin (CAS: 385437-57-0), and (Δ9-Cis) PC (DOPC) (CAS: 4235-95-4) were purchesed from Avanti Polar Lipids INC, ATTO 655-DOPE (order code: AD 655-161) and ATTO 488 carboxy (order code: AD 488-21) were purchased from ATTO-TEC. For the single particle assay eGFPsiRNA_sense3'AlexF647N was used and purchased from Integrated DNA Technologies. For the eGFP transfection experiments in HEK293 cells stably expressing d2eGFP, Silencer GFP (eGFP) siRNA was used and purchesed from Ambion. pLV-CMV-d2eGFP plasmid (Vector ID: VB900138-0467pjn) was ordered from VectorBuilder.

#### Substrate surface preparation for single particle imaging

Glass slides 76 x 26 nm were dried with a nitrogen flow and activated using a plasma cleaner for 1 minute at a pressure between 300 mtorr to 400 mtorr. Immediately after activation the slides were attached to the glass slides in order to make flow cells. Each of the six champers was passivated with 80 μL of a PLL-g-PEG (1 mg/mL in HEPES buffer pH 7.8) and PLL-g-PEG-biotin (1 mg/mL in HEPES buffer pH 5.5) mixture in the ratio 100:1. The surface was incubated for 30 min. Excess of PLL-g-PEG mixture was removed by flushing five times with 50 μL HEPES buffer pH 7.8. 80 μL of the neutravidin solution (1 mg/mL in HEPES buffer pH 7.8) was added in each well. The surface was incubated for 15 min. Each well was washed with 10 mM PBS buffer pH 7.6 at least 5 times before the sample addition.

#### Bovine Serum Albumin (BSA) labelling with ATTO 655

Native BSA was labelled with ATTO 655 NHS-ester on free lysine residues by mixing a 20 mg/mL ATTO 655 NHS-ester solution in DMSO (45 μL, 1.1 x $10^{-3}$ mmol, 30.0 equiv.) with a 300 uM BSA solution in 10 mM PBS buffer pH 8.4 (2.3 mg, 3.5 x $10^{-5}$ mmol, 115 uL, 1.0 equiv.) and dilute in 1.0 mL final volume (35 uM of the BSA in the final solution) for 2 hours at room temperature. Free dye were removed by dialysis using an 8-10 kDa MWCO membrane against 10 mM PBS buffer pH 8.4 (3 x 2 hours). Labelling efficiency was characterized by Nanodrop and found to be 154 $\pm$ 4 % (as more than one lysine residues are present on the surface of BSA). Labelled BSA was stored in 35 uM aliquots of 15 μL at -20 °C. Each aliquot was only used once.

### 2. Analytical Techniques

#### Id-3 spectraMax plate reader

Liposome storage stability and MTS-PMS cell death experiments were performed using an Id-3 spectraMax plate reader. Clear standard 96-well plates were used to measure absorbance, while black standard 96-well plates were used to measure fluorescence, to minimize interference between wells.

#### DLS measurements

The liposomes size distribution and response studies were determined by dynamic light scattering using a Malvern Zetasizer μV apparatus (Malvern Panalytical, UK) following the manufacturer's instructions. The sample was diluted to reach a final concentration of 0.005 mg/mL, filtered and its polydispersity index was measured at 25 or 37 °C. For the Refractive Index, LNPs was chosen. A QS High Precision Cell Cuvette made of Quartz SUPRASIL was used (Art. No. 105-231-001-8.5-40, light pate 1.25x1.25, center 8.5).

**Nanodrop**

A Thermo Scientific™ Invitrogen™ Nanodrop™ One Spectrophotometer with Qubit™ 4 Fluorometer was used for the labelling efficiency calculation of the labelled Bovine Serum Albumin (BSA).

**Total internal reflection fluorescence (TIRF) microscopy**

A TIRF microscope (IX83, Olympus) was used for the single-particle tracking (SPT) experiments. Oil immersion objective (UAPON 100XOTIRF, NA 1.49, Olympus) and an EMCCD camera (ImagEM X2, Hamamatsu, Shizuoka, Japan) were used to record images and videos with a pixel width of 160 nm and field of view 81.92 μm × 81.92 μm. Laser lines of 640 nm and 488 nm were used to excite the fluorophores ATTO-655-DOPE (liposome surface) and ATTO-488 carboxy (cargo), respectively. Imaging was performed with an exposure time of 50 ms, 100 nm penetration depth, and 300 EM gain. Each image series with a frame rate 2 frames per 5 seconds, contained 10 frames of the 488 nm channel (1.3 mW) and 10 frames of the red channel (0.9 mW) alternating between them. Prior to imaging the sample (C = 1.5 $10^{-2}$ mg/mL) was incubated for 10 minutes and washed with 10 mM PBS buffer pH 7.6 (6 x 0.5 mL).

**Cell culture**

HeLa and HEK293 cells were cultured in Nunc EasYFlask 25 cm2 cell culture flasks with Dulbecco's Modified Eagle's Medium with 4500 mg/L glucose supplemented with 2 mM L-glutamine, 1 mM sodium pyruvate, and 10% (v/v) heat-inactivated FBS. hCMEC/D3 cells were cultured in EBM-2 endothelial basal medium (Lonza) supplemented with 2mM L-glutamine and 10% (v/v) heat-inactivated FBS. Both cell lines were detached by 3 × trypsin-EDTA treatment (5 mg/mL trypsin and 2 mg/mL EDTA in DPBS, pH 7.4) and seeded for subsequent experiments. Cells were incubated in a humidified, 5% CO2, 37oC incubator. The passages for all cell experiments ranged from 7 to 14.

For MTS-PMS assay: Cells were seeded at a density of 2.5 × 104 cells/well in 96-well plates containing Dulbecco's Modified for cell viability assessment, 24 h before the experiments.

**Spinning Disk Confocal Microscopy (SDCM)**

An oil immersion 60x objective (Olympus) and a numerical aperture of 1.4 connected to a CMOS camera (photometric PRIME 95B) with an effective pixel size of 183 nm x 183 nm was used for cell imaging. Cells were grown into 8-well plates (Ibidi) at a density of 15.000 cells per well containing 200 μL Dulbecco's Modified Eagle's Medium.

*Particle internalization studies*: After incubation, the culture media was removed, and the cells were washed twice with PBS. Subsequently, 200 μL of imaging media and 0.4 μL of ATTO 488 Carboxyl 50 mM dye exclusion were added to each well, followed by the addition of 1 μL of liposome (membrane-labelled with ATTO 655) solution of a concentration of 1 mg/mL. For each experimental condition, 10 positions were imaged in parallel directly after the addition of the liposomes and every 1h in a 3h-time period. Imaging was performed with an exposure time of 50.04 ms. Each image series with a frame rate 2 frames per 1 seconds contained 100 frames of the 488 nm channel and then 100 frames of the 647 nm channel.

*eGFP Transfection studies*: HEK293 cells stably expressing d2eGFP were established by transducing cells with a lentiviral vector encoding d2eGFP. The lentiviral particles were produced by transfecting the HEK293T packaging cell line with pMD2.G, psPAX2, and pLV-CMV-d2eGFP plasmid using Lipofectamine 3000 according to the manufacturer's protocol. HEK293 cells stably expressing d2eGFP were transferred to a preheated microscopy incubation chamber, and 5 positions with evenly distributed cells were selected for each experimental condition (cells in the absence of particles, cells treated with siRNA loaded 5% poly(DPA) modified liposomes, and cells treated with siRNA loaded non-modified liposomes). Immediately before starting image acquisition, of 1 µL of liposome solution of a concentration of 1 mg/mL loaded with siRNA targeting d2eGFP were added dropwise to the medium of the wells of interest. Seven z-plane images with 3 µm z-spacing were acquired per position at 2.5 min intervals. Images were acquired for 10 h in total. For the excitation of the d2eGFP, a blue laser (488nm) was used with a 50,04 ms exposure time.

### 3. Functionalized liposome preparation

All liposome populations were prepared following a well-established method.[1] The desired mole percentage of lipids was mixed in a glass vial using glass syringes. The syringes were cleaned before and between switching between lipids 20 times with chloroform. The glass vial was dried by softly flowing pure nitrogen for 10 -15 minutes until the lipids formed a thin film. The vial was furthermore vacuumed for at least one hour. The lipids were rehydrated to a liposome concentration of $1 gL^{-1}$ with 10 mM PBS buffer pH 7.6 and vortexed for 30 seconds. The liposomes samples were incubated for 60 minutes to self-assemble and then were flash-frozen in a mixture of dry ice and acetone followed by thawing in a heat bath. This was repeated 10 times to ensure unilamellarity. To ensure homogeneous size distribution, the samples were extruded 10 times (back and forth) through a 100 nm pore size membrane. The samples were aliquoted and stored at -20 °C.

**Lipid composition for non-modified liposome**: HSPC 64%, DOPC 20%, Cholesterol 15%, DSPE-PEG-biotin 0.5%, ATTO-655-DOPE 0.5%. The lipids were rehydrated with 470 µL PBS buffer 10 mM pH 7.6 mixed with 30 µL ATTO-488 carboxy (500 µM in DMSO).

**Lipid composition for empty non-modified liposome (control experiments)**: HSPC 64%, DOPC 20%, Cholesterol 15%, DSPE-PEG-biotin 0.5%, ATTO-655-DOPE 0.5%. The lipids were rehydrated with 500 µL PBS buffer 10 mM pH 7.6.

**Lipid composition for 5% poly(DPA) modified liposome**: HSPC 59%, DOPC 20%, Cholesterol 15%, DSPE-PEG-biotin 0.5%, ATTO-655-DOPE 0.5%, poly(DPA) 5%. The lipids were rehydrated with 470 µL PBS buffer 10 mM pH 7.6 mixed with 30 µL ATTO-488 carboxy (500 µM in DMSO).

**Lipid composition for empty 5% poly(DPA) modified liposome (control experiments)**: HSPC 59%, DOPC 20%, Cholesterol 15%, DSPE-PEG-biotin 0.5%, ATTO-655-DOPE 0.5%, poly(DPA) 5%. The lipids were rehydrated with 500 µL PBS buffer 10 mM pH 7.6.

**Lipid composition for 1% poly(DPA) modified liposome**: HSPC 63%, DOPC 20%, Cholesterol 15%, DSPE-PEG-biotin 0.5%, ATTO-655-DOPE 0.5%, poly(DPA) 1%. The lipids were rehydrated with 470 µL PBS buffer 10 mM pH 7.6 mixed with 30 µL ATTO-488 carboxy (500 µM in DMSO).

**Lipid composition for empty 1% poly(DPA) modified liposome (control experiments)**: HSPC 59%, DOPC 20%, Cholesterol 15%, DSPE-PEG-biotin 0.5%, ATTO-655-DOPE 0.5%, poly(DPA) 5%. The lipids were rehydrated with 500 µL PBS buffer 10 mM pH 7.6.

**Lipid composition for 1% poly(DEAEMA) modified liposome**: HSPC 63%, DOPC 20%, Cholesterol 15%, DSPE-PEG-biotin 0.5%, ATTO-655-DOPE 0.5%, poly(DEAEMA) 1%. The lipids were rehydrated with 470 μL PBS buffer 10 mM pH 7.6 mixed with 30 μL ATTO-488 carboxy (500 μM in DMSO).

**Lipid composition for empty 1% poly(DEAEMA) modified liposome (control experiments)**: HSPC 63%, DOPC 20%, Cholesterol 15%, DSPE-PEG-biotin 0.5%, ATTO-655-DOPE 0.5%, poly(DEAEMA) 1%. The lipids were rehydrated with 500 μL PBS buffer 10 mM pH 7.6.

**Lipid composition for 5% poly(DPA) modified liposome loaded with methylene blue**: HSPC 59%, DOPC 20%, Cholesterol 15%, DSPE-PEG-biotin 0.5%, ATTO-655-DOPE 0.5%, poly(DPA) 5%. The lipids were rehydrated with 470 μL PBS buffer 10 mM pH 7.6 mixed with 30 μL methylene blue (500 μM in DMSO).

**Lipid composition for 5% poly(DPA) modified liposome loaded with siRNA**: HSPC 59%, DOPC 20%, Cholesterol 15%, DSPE-PEG-biotin 0.5%, ATTO-655-DOPE 0.5%, poly(DPA) 5%. The lipids were rehydrated with 190 μL PBS buffer 10 mM pH 7.6 mixed with 10 μL siRNA (10 μM in PBS buffer). The freeze-thaw cycle step was skipped to avoid siRNA degradation.

**Lipid composition for 5% poly(DPA) modified liposome loaded with BSA**: HSPC 59%, DOPC 20%, Cholesterol 15%, DSPE-PEG-biotin 0.5%, ATTO-655-DOPE 0.5%, poly(DPA) 5%. The lipids were rehydrated with 500 μL of BSA solution in PBS buffer 10 mM pH 7.6 (30 μM in PBS buffer). The freeze-thaw cycle step was skipped to avoid enzyme denaturation.

### 4. Quantification of the liposome population's encapsulation efficiency (EE)

All the liposome populations were labelled on their membrane with ATTO-655 fluorophore (detected in the red channel), whereas they were loaded with ATTO-488 carboxy (detected in the blue channel). The dual color imaging on TIRF allows for the encapsulation efficiency of each liposome population to be defined as EE = $N_o$ of Loaded Liposomes/$N_o$ of Total Liposomes. For the intensity extractions, an in-house, GPU accelerated, Laplacian of Gaussian was used for particle detection and intensity extraction. Nearest Neighbours was implemented on the GPU accelerated framework for particle tracking. Custom Python scripts were developed and used for the post-detection intensity analysis and statistics for both static and temporal experiments. The detection with LoG was performed in the liposome membrane channel. For each one of the detections with position (xi, yi) and radius of σi we can perform the intensity integration and background correction in the cargo channel on the respective positions and ROIs. Doing so and tracking the detected particles while integrating their intensities in the cargo channel we defined the loaded particles. The same liposome populations were prepared without loading ATTO-488 carboxy. The extracted ATTO-488 carboxy signal was compared to the signal of non-loaded liposomes as a control experiment to ensure that the signal can be attributed to loaded ATTO-488 carboxy and not to the crosstalk between the two channels, setting a threshold of 95% confidence. The laser line of 640 nm was used to excite ATTO-655, and 488 nm was used to excite the loaded ATTO-488 carboxy. Movies of 10 frames of the liposomes in the red and in blue channel (ATTO-488 carboxy channel) were recorded alternating between them. For 5% poly(DPA) modified liposomes loaded with methylene blue, siRNA and BSA protein the liposome membrane was labelled with ATTO-488 (detected on the blue channel). Methylene blue has emission max at 688 nm (detected on the red channel), siRNA was labelled with AlexF647N detected on the red channel and BSA was labelled with ATTO-655 NHS ester lebelled on the red channel. The laser line of 488 nm was used to excite the liposome membrane and the laser line of 640 nm was used to excite each of the cargos.

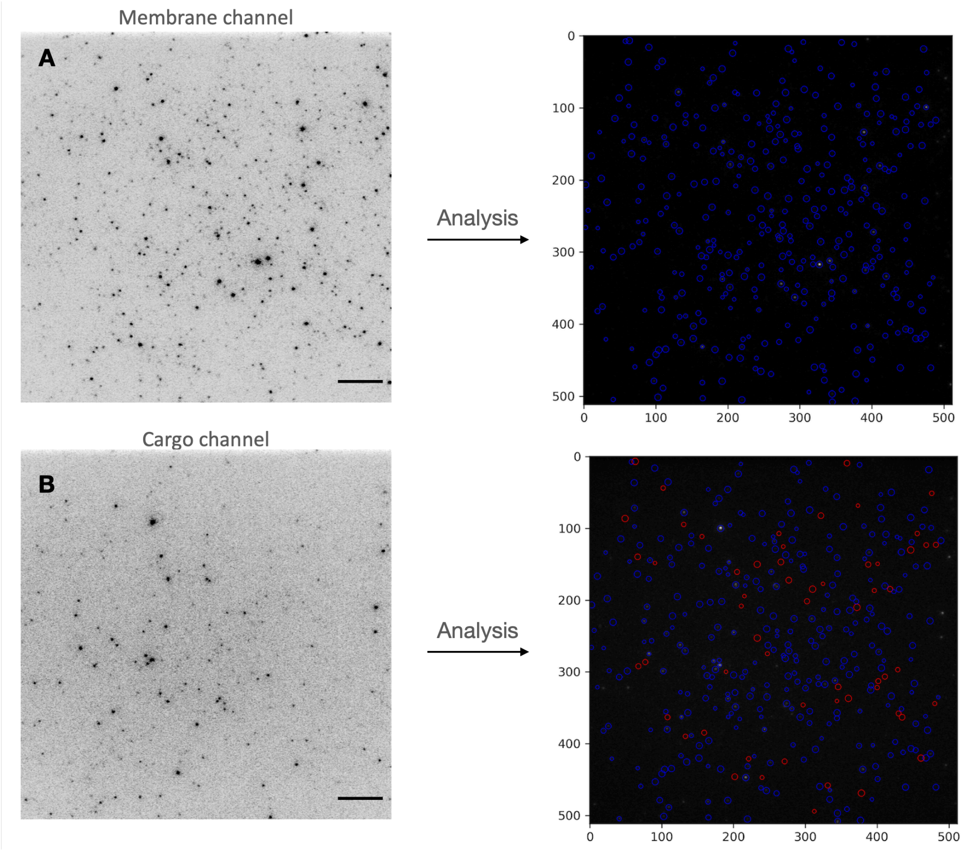


| Liposome Modification | Cargo Type | Total Nₒ of particles | Total Nₒ of loaded particles | Total Nₒ of empty particles | Encapsulation efficiency (%) |
|---|---|---|---|---|---|
| **Non-modified** | ATTO-488 | 7772.0 | 4106.0 | 3666.0 | 53 |
| **5% poly(DPA)** | ATTO-488 | 4362.0 | 2857.0 | 1505.0 | 65 |
| **1% poly(DPA)** | ATTO-488 | 7201.0 | 2440.0 | 4761.0 | 34 |
| **1% poly(DEAEMA)** | ATTO-488 | 8486.0 | 1065.0 | 7421.0 | 13 |
| **5% poly(DPA)** | Methylene Blue | 4388.0 | 3126.0 | 1262.0 | 71 |
| **5% poly(DPA)** | siRNA | 51484.0 | 23579.0 | 27905.0 | 55 |
| **5% poly(DPA)** | BSA | 24726.0 | 9820.0 | 14906.0 | 40 |

**Figure S1**. **A**. Left: Representative TIRF images of a single field of view imaging thousands of surface-tethered polymer modified liposome at pH 7.6 on the membrane channel. Scale bar corresponds to 1 um. Right: Particle detection in the liposome membrane channel using an in-house, GPU accelerated, Laplacian of Gaussian. **B**. Left: Representative TIRF images of a single field of view imaging thousands of surface-tethered polymer modified liposome at pH 7.6 on the cargo channel. Scale bar corresponds to 1 um. Right: Particle detection in the cargo channel using an in-house, GPU accelerated, Laplacian of Gaussian. Custom Python scripts were developed and used for the post-detection intensity analysis and applying a theshold of 96% confidence the empty and non empty particles were distinguished. Blue: non empty particles. Red: empty particles. **Table**: Overall encapsulation efficiency values for all the prepared liposome populations. The mean values correspond to the mean of Nₒ of Loaded Liposomes/Total Nₒ of Liposomes of six fields of view. Error bars corespond to the standard deviation of the mean values of six technical replicates each.

**Encapsulation efficiency per size**

The empty particles of each liposome populations were categorized in size groups using a bin size of 40 nm The percentage of empty particles in each size group was quantified.

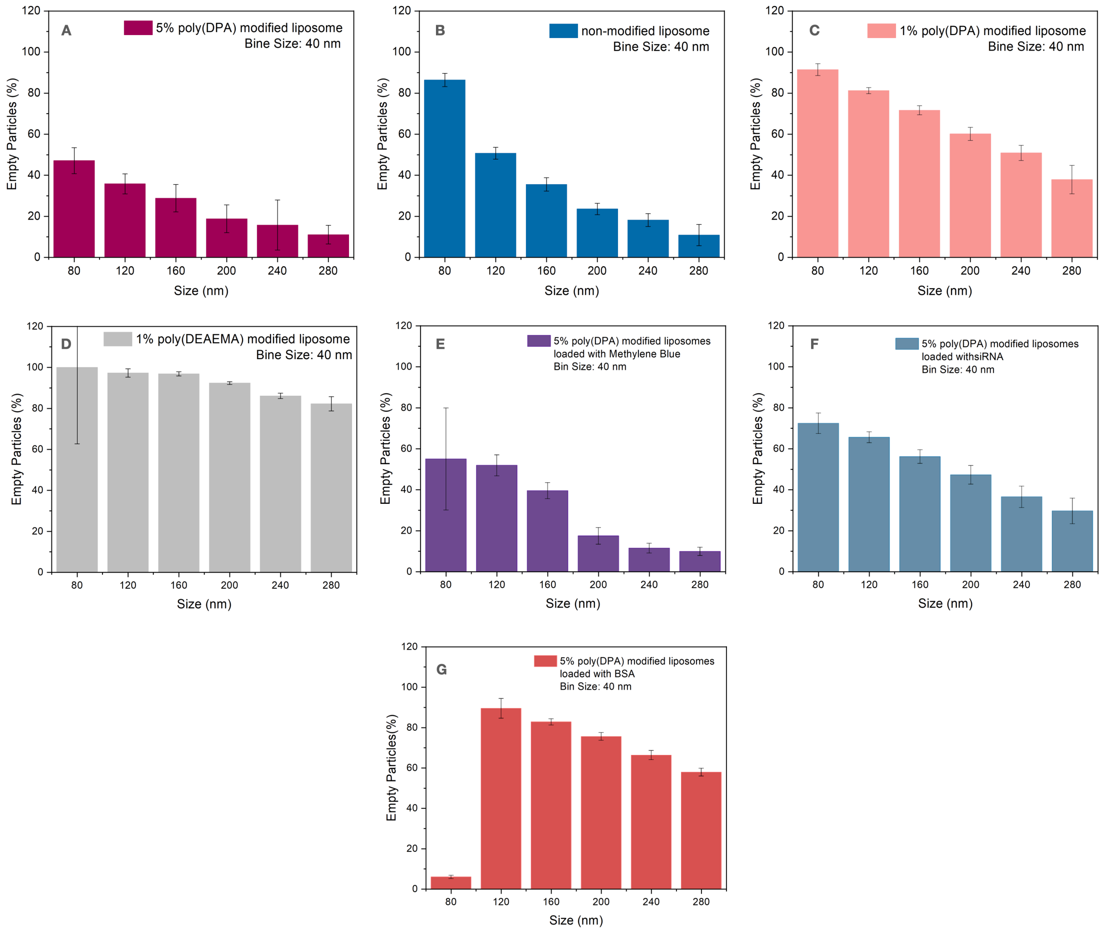


**Figure S2**. Percentage of empty particles at 6 different size groups using 40 nm binning. **A**. Liposome population modified with 5% poly(DPA) loaded with ATTO-488 carboxy. **B**. Non-modified liposome population loaded with ATTO-488 carboxy. **C**. Liposome population modified with 1% poly(DPA) loaded with ATTO-488 carboxy. **D**. Liposome population modified with 1% poly(DEAEMA) loaded with ATTO-488 carboxy. **E**. Liposome population modified with 5% poly(DPA) loaded with methylene blue. **F**. Liposome population modified with 5% poly(DPA) loaded with siRNA. **G**. Liposome population modified with 5% poly(DPA) loaded with BSA protein. Error bars corespond to the standard deviation of the mean values of six technical replicates.

Further single particle analysis with 40 nm binning revealed sizer-dependent effects on loading profiles. Cargo intensity scaled with size across all liposome population, as expected, but when normalized to vesicle volume, smaller vesicles demonstrate significantly higher cargo density indicating more efficient packing.

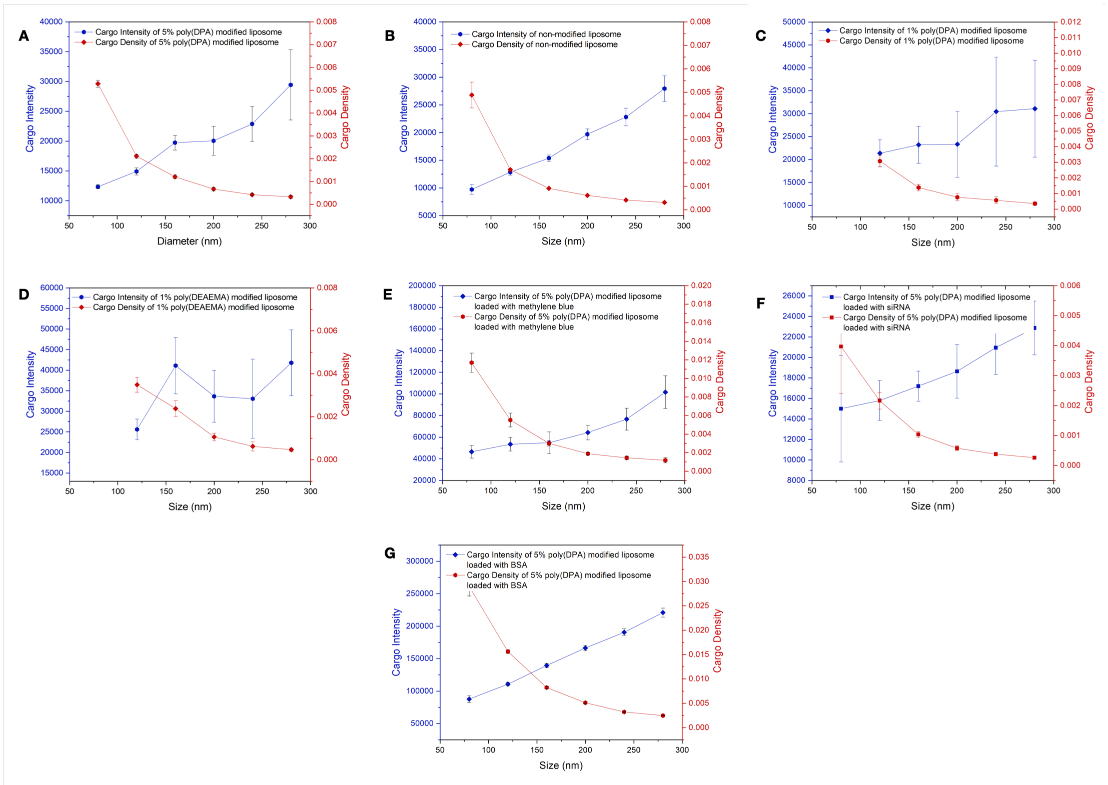


**Figure S3**. **A**. Cargo intensity versus cargo density for the 5% poly(DPA) modified liposomes loaded with ATTO-488 carboxy. **B**. Cargo intensity versus cargo density for the non-modified liposomes loaded with ATTO-488 carboxy. **C**. Cargo intensity versus cargo density for the 1% poly(DPA) modified liposomes loaded with ATTO-488 carboxy. **D**. Cargo intensity versus cargo density for the 1% poly(DEAEMA) modified liposomes loaded with ATTO-488 carboxy. **E**. Cargo intensity versus cargo density for the 5% poly(DPA) modified liposomes loaded with methylene blue. **F**. Cargo intensity versus cargo density for the 5% poly(DPA) modified liposomes loaded with siRNA. **G**. Cargo intensity versus cargo density for the 5% poly(DPA) modified liposomes loaded with BSA. The bin size is 40 nm. Error bars correspond to the standard deviation of the median intensity of six replicates.

### 5. Effect of pH on cargo release

All the liposomes were loaded with ATTO-488 carboxy (30 μM), and the pH triggered cargo release, and the loaded particle loss were evaluated upon imaging these liposomes at decreasing pH conditions with total internal reflection (TIRF) microscopy. For this purpose, six-champer glass slides functionalized with neutravidin were used for liposome immobilization at 10 mM PBS pH 7.6 (313 mOsm/L) and then the buffer solution was exchanged to 156 mM sodium acetate buffer pH 5.2 (313 mOsm/L). Each well was washed at least 5 times before every measurement with the buffer solution of the chamber in order for the fluorophore that is released in the solution to be removed. The laser line of 488 nm was used to excite the loaded ATTO-488 carboxy. Movies of 10 frames of the liposomes in the blue channel (ATTO-488 carboxy channel) were recorded. The intensity histograms are fitted with the LogNormal fit. The mu values and the sigma of the LogNormal fit are displayed.

#### 5.1. Effect of pH on cargo release

Image series of 6 fields of view were recorded in the blue channel (where ATTO-488 is detected) at pH 7.6 and at pH 5.2 at the following time points: 0 min, 30 min, 60 min and 120 min. For the modified liposomes Image series of 6 fields of view were also recorded on the blue channel at pH 7.6 at the following time points: 0 min, 30 min, 60 min, and 120 min to evaluate if the intensity loss corresponds to ATTO-488 carboxy photobleaching.

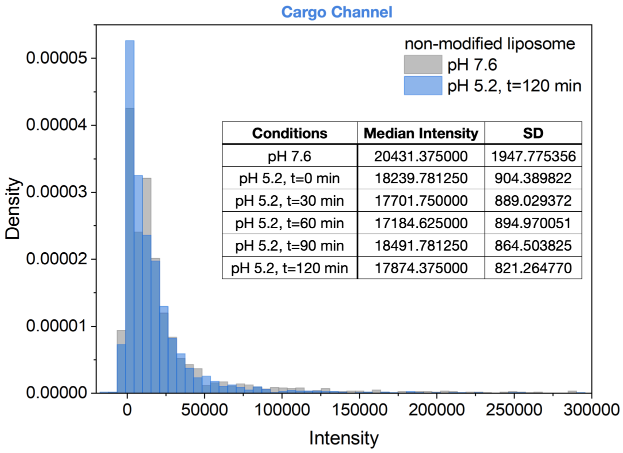


| Conditions | Median Intensity | SD |
|---|---|---|
| pH 7.6 | 20431.375000 | 1947.775356 |
| pH 5.2, t=0 min | 18239.781250 | 904.389822 |
| pH 5.2, t=30 min | 17701.750000 | 889.029372 |
| pH 5.2, t=60 min | 17184.625000 | 894.970051 |
| pH 5.2, t=90 min | 18491.781250 | 864.503825 |
| pH 5.2, t=120 min | 17874.375000 | 821.264770 |

**Figure S4**. Median intensity values of non-modified liposome loaded with ATTO-488 in the blue channel at pH 7.6 and at pH 5.2 immediately after pH drop and after 30 min, 60 min, and 120 min detected at six fields of view. Grey: Distribution of background corrected intensities of non-modified liposome in the blue channel at pH 7.6. Blue: Distribution of background corrected intensities of non-modified liposome in the blue channel at pH 5.2 after 120 min of the pH drop. Insert table: median intensity values at pH 7.6 and pH 5.2 over time. Error bars correspond to the standard deviation of six fields of view.

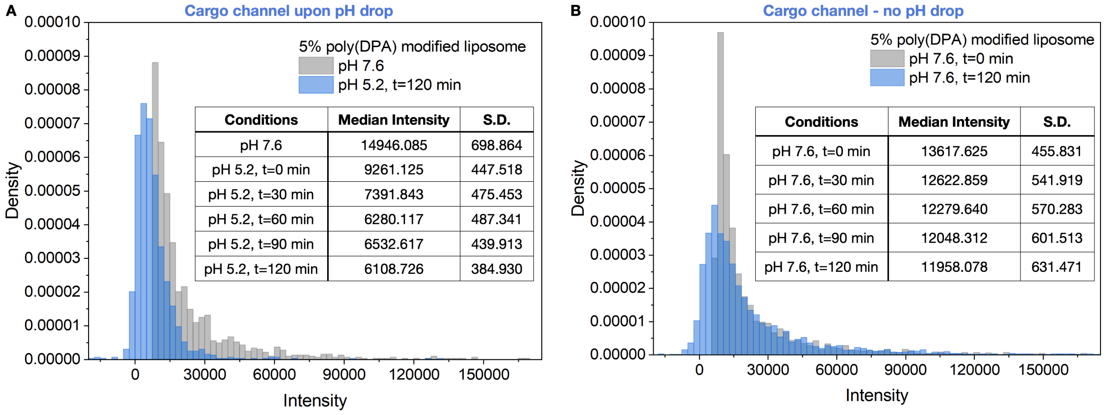


| Conditions | Median Intensity | S.D. |
|---|---|---|
| pH 7.6 | 14946.085 | 698.864 |
| pH 5.2, t=0 min | 9261.125 | 447.518 |
| pH 5.2, t=30 min | 7391.843 | 475.453 |
| pH 5.2, t=60 min | 6280.117 | 487.341 |
| pH 5.2, t=90 min | 6532.617 | 439.913 |
| pH 5.2, t=120 min | 6108.726 | 384.930 |

| Conditions | Median Intensity | S.D. |
|---|---|---|
| pH 7.6, t=0 min | 13617.625 | 455.831 |
| pH 7.6, t=30 min | 12622.859 | 541.919 |
| pH 7.6, t=60 min | 12279.640 | 570.283 |
| pH 7.6, t=90 min | 12048.312 | 601.513 |
| pH 7.6, t=120 min | 11958.078 | 631.471 |

**Figure S5**. Median intensity values of 5% poly(DPA) modified liposome loaded with ATTO-488 in the blue channel at pH 7.6 and at pH 5.2 immediately after pH drop and after 30 min, 60 min, and 120 min detected at six fields of view. Grey: Distribution of background corrected intensities of 5% poly(DPA) modified liposome in the blue channel at pH 7.6. Blue: Distribution of background corrected intensities of 5% poly(DPA) modified liposome in the blue channel at pH 5.2 after 120 min of the pH drop. Insert table: median intensity values at pH 7.6 and pH 5.2 over time. Error bars correspond to the standard deviation of six fields of view.

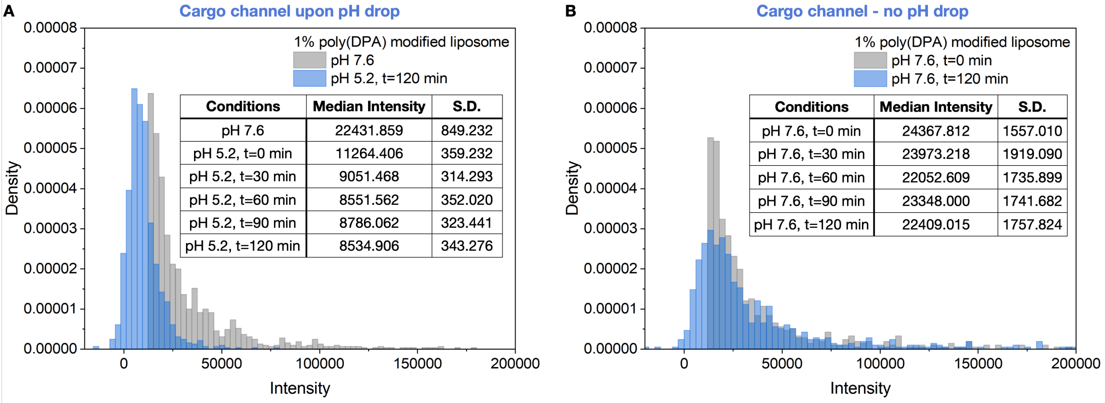


**Figure S6**. Median intensity values of 1% poly(DPA) modified liposome loaded with ATTO-488 in the blue channel at pH 7.6 and at pH 5.2 immediately after pH drop and after 30 min, 60 min, and 120 min detected at six fields of view. Grey: Distribution of background corrected intensities of 1% poly(DPA) modified liposome in the blue channel at pH 7.6. Blue: Distribution of background corrected intensities of 1% poly(DPA) modified liposome in the blue channel at pH 5.2 after 120 min of the pH drop. Insert table: median intensity values at pH 7.6 and pH 5.2 over time. Error bars correspond to the standard deviation of six fields of view.

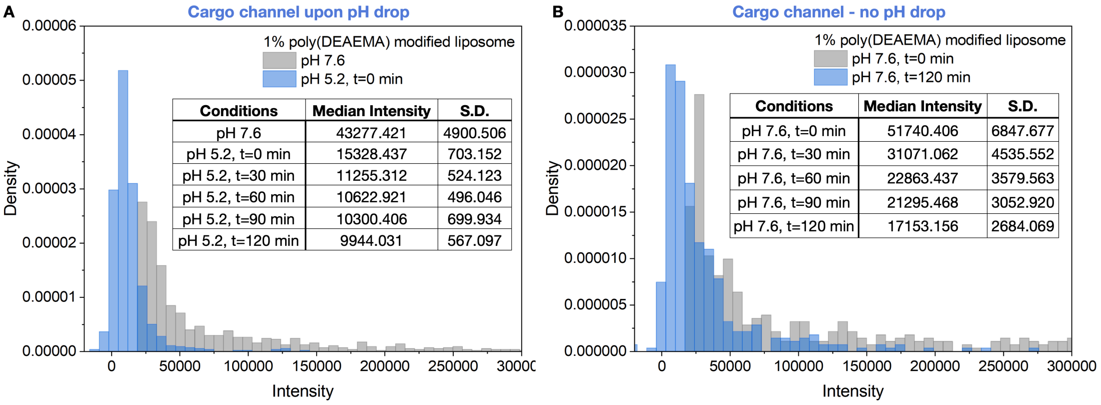


**Figure S7**. Median intensity values of 1% poly(DEAEMA) modified liposome loaded with ATTO-488 in the blue channel at pH 7.6 and at pH 5.2 immediately after pH drop and after 30 min, 60 min, and 120 min detected at six fields of view. Grey: Distribution of background corrected intensities of 1% poly(DEAEMA) modified liposome in the blue channel at pH 7.6. Blue: Distribution of background corrected intensities of 1% poly(DEAEMA) modified liposome in the blue channel at pH 5.2 after 120 min of the pH drop. Insert table: median intensity values at pH 7.6 and pH 5.2 over time. Error bars correspond to the standard deviation of six fields of view.

#### Cargo release per size

Using 40 nm size binning, cargo release was correlated to particle size, revealing that all particle sizes respond to pH drop and release their cargo.

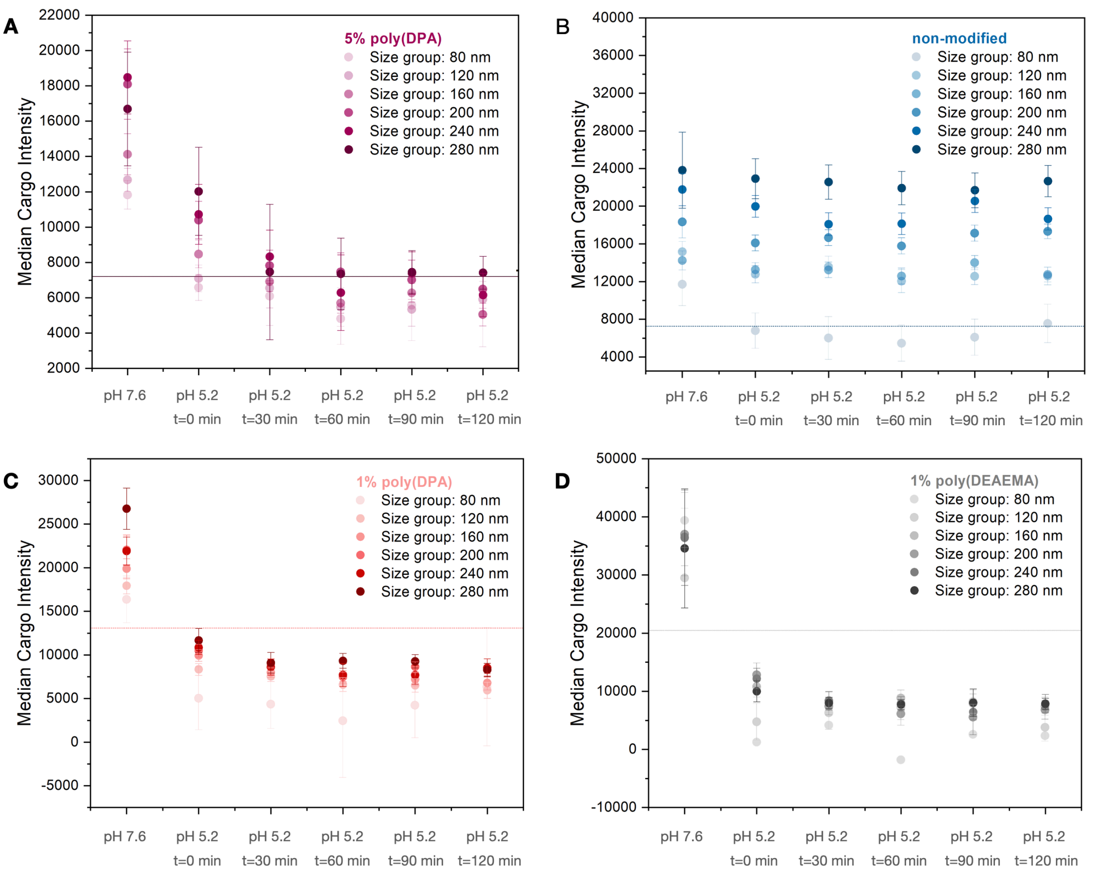


**Figure S8**. **A**. Median cargo intensity values of the 5% poly(DPA) modified liposome at pH 7.6 and at pH 5.2 at 5 time points from 0 to 120 min per size. Data binned every 40 nm starting from 80 nm. Error bars correspond to the standard deviation of the median intensities of two biological replicates. The threshold at 7206 distinguishes the empty and the non-empty particles. **B**. Median cargo intensity values of the non-modified liposome at pH 7.6 and at pH 5.2 at 5 time points from 0 to 120 min per size. Data binned every 40 nm starting from 80 nm. Error bars correspond to the standard deviation of the median intensities of two biological replicates. The threshold at 7278 distinguishes the empty and the non-empty particles. **C**. Median cargo intensity values of the 1% poly(DPA) modified liposome at pH 7.6 and at pH 5.2 at 5 time points from 0 to 120 min per size. Data binned every 40 nm starting from 80 nm. Error bars correspond to the standard deviation of the median intensities of two biological replicates. The threshold at 13084 distinguishes the empty and the non-empty particles. **D**. Median cargo intensity values of the 1% poly(DEAEMA) modified liposome at pH 7.6 and at pH 5.2 at 5 time points from 0 to 120 min per size. Data binned every 40 nm starting from 80 nm. Error bars correspond to the standard deviation of the median intensities of two biological replicates. The threshold at 20477 distinguishes the empty and the non-empty particles.

### Effect of pH on ATTO-488 carboxy fluorescence

The effect of pH on ATTO-488 carboxy fluorescence was evaluated to ensure that the intensity loss in the blue channel is attributed to ATTO-488 carboxy release and not to changes on the fluorescent properties of the fluorophore. The fluorescence of ATTO-488 carboxy at 520 nm was measured under our experimental pH conditions. A small fluorescence increase was observed by dropping the pH from 7.6 to 5.2 indicating that the intensity loss in the blue channel does not correspond to changes on ATTO-488 carboxy fluorescence.

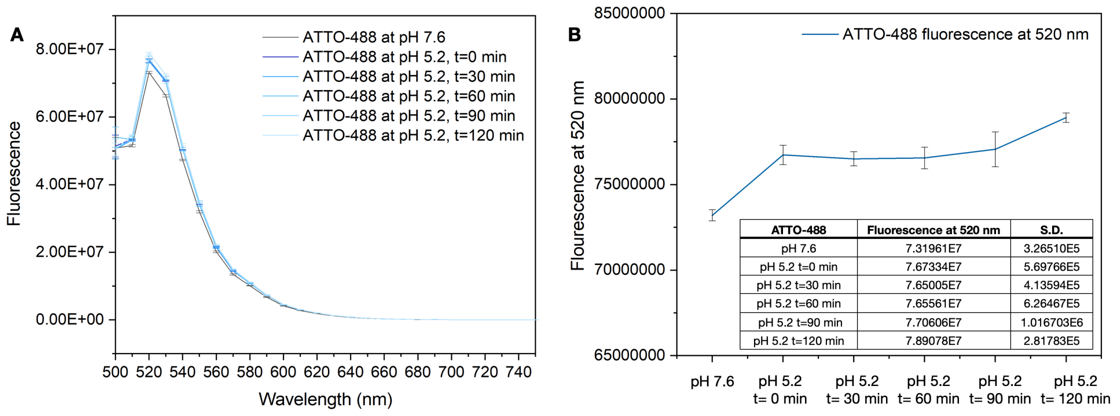


**Figure S9**. **A**. Fluorescence spectra of 100 nM solution ATTO-488 carboxy in PBS pH 7.6 and acetate buffer pH 5.2 immediately after the pH drop and after 30, 60, 90 and 120 min. **B**. Fluorescence values and the standard deviation of the values at 520 nm. The measurements were conducted in triplicates.

### 6. Effect of pH on liposome membrane

The effect of pH on the membrane of the non-polymer modified liposome and 5% pH responsive poly(DPA) was evaluated. The liposome membrane was decorated with ATTO-655 intensity of the membrane was evaluated upon imaging these liposomes at decreasing pH conditions with total internal reflection (TIRF) microscopy. For this purpose, six-champer glass slides functionalized with neutravidin were used for liposome immobilization at 10 mM PBS pH 7.6 (313 mOsm/L) and then the buffer solution was exchanged to 156 mM sodium acetate buffer pH 5.2 (313 mOsm/L). Each well was washed at least 5 times before every measurement with the buffer solution of the chamber in order for the fluorophore that is released in the solution to be removed. The laser line of 640 nm was used to excite ATTO-655. Movies of 10 frames of the liposomes in the red channel (ATTO-655 channel) were recorded. The intensity histograms are fitted with the LogNormal fit. The mu values and the sigma of the LogNormal fit are displayed.
Image series of 6 fields of view were recorded in the red channel (where ATTO-655 is detected) at pH 7.6 and at pH 5.2 at the following time points: 0 min, 30 min, 60 min, and 120 min. For the pH responsive liposome Image series of 6 fields of view were also recorded on the red channel at pH 7.6 at the following time points: 0 min, 30 min, 60 min, and 120 min to evaluate if the intensity loss corresponds to ATTO-655 photobleaching.

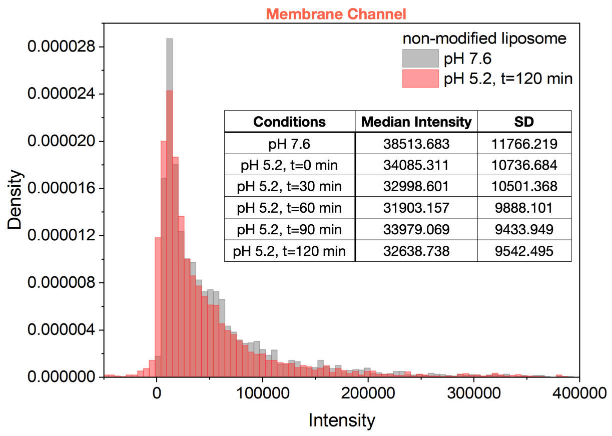


**Figure S10**. **A**. Median intensity values of non-modified liposome decorated with ATTO-655 in the red channel at pH 7.6 and at pH 5.2 immediately after pH drop and after 30 min, 60 min, 90 min and 120 min detected at six fields of view. **B**. Grey: Distribution of background corrected intensities of liposome 4 in the red channel at pH 7.6. Red: Distribution of background corrected intensities of liposome 4 in the red channel at pH 5.2 immediately after the pH drop. Red: Distribution of background corrected intensities of liposome 4 in the red channel at pH 5.2 after 1200 min of the pH drop. Insert table: median intensity values at pH 7.6 and pH 5.2 over time. Error bars correspond to the standard deviation of six fields of view.

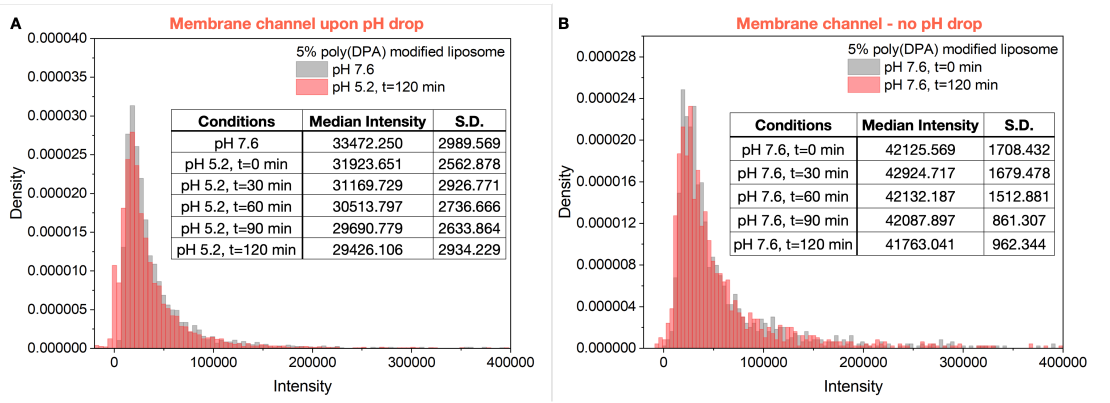


**Figure S11**. **A**. Median intensity values of liposome 3 decorated with ATTO-655 in the red channel at pH 7.6 and at pH 5.2 immediately after pH drop and after 30 min, 60 min, 90 min and 120 min detected at six fields of view. **B**. Grey: Distribution of background corrected intensities of liposome 3 in the red channel at pH 7.6. Red: Distribution of background corrected intensities of liposome 3 in the red channel at pH 5.2 immediately after the pH drop. Red: Distribution of background corrected intensities of liposome 3 in the red channel at pH 5.2 after 90 min of the pH drop. Insert table: median intensity values at pH 7.6 and pH 5.2 over time. Error bars correspond to the standard deviation of six fields of view.

#### Effect of pH on ATTO-655 fluorescence

The effect of pH on ATTO-655 fluorescence was evaluated to evaluate the pH triggered changes on the fluorescent properties of the fluorophore. The fluorescence of ATTO-655 at 680 nm was measured under our experimental pH conditions. A small fluorescence decrease of 5% was observed by dropping the pH from 7.6 to 5.2 indicating that the pH drop can result in a small intensity loss in the red channel.

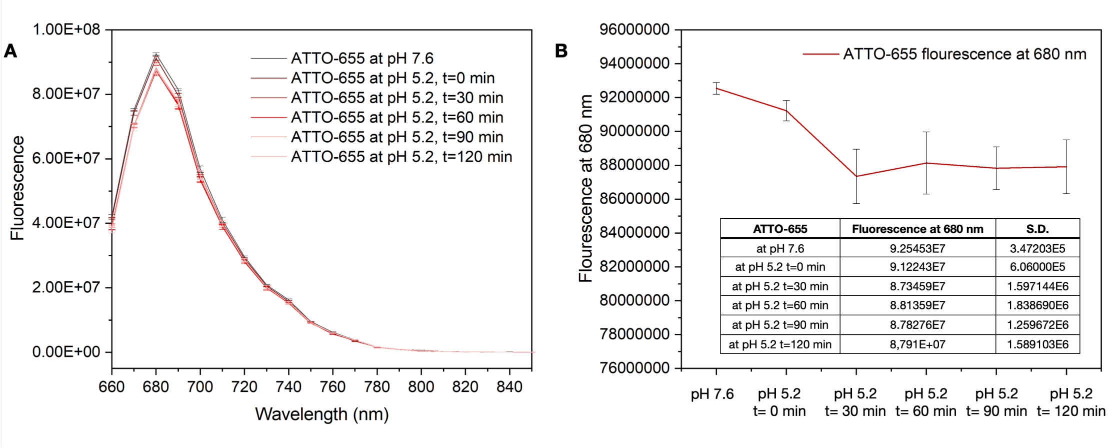


**Figure S12**. **A**. Fluorescence spectra of 5 μM solution ATTO-655 in PBS pH 7.6 and acetate buffer pH 5.2 immediately after the pH drop and after 30, 60, 90 and 120 min. **B**. Fluorescence values and the standard deviation of the values at 680 nm. The measurements were conducted in triplicates.

### 7. Non-modified and 5% poly(DPA) modified liposome leakage over time

#### Calibration curve of ATTO-488 carboxy (cargo)

The maximum absorbance of ATTO-488 carboxy at 500 nm was measured using ATTO-488 carboxy solutions of 10$^{-2}$ μmol/mL, 8 10$^{-3}$ μmol/mL, 6 10$^{-3}$ μmol/mL, 4 10$^{-3}$ μmol/mL, 2 10$^{-3}$ μmol/mL, 10$^{-3}$ μmol/mL and 5 10$^{-4}$ μmol/mL in PBS pH 7.6. The formula for linear regression was **f(x)= 130.99744x + 0.03303**.

A

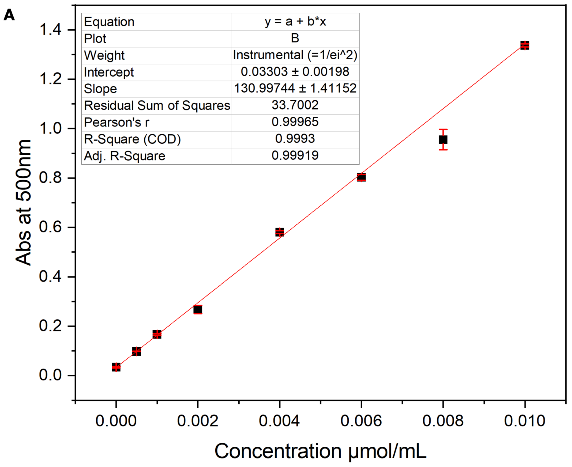


B

| Concentration μmol/mL | Absorbance at 500 nm |
|---|---|
| 0 | 0.03393 +/- 0.0025 |
| 0,01 | 1.33703 +/- 0.0064 |
| 0,008 | 0.9557 +/- 0.04133 |
| 0,006 | 0.80257 +/- 0.01239 |
| 0,004 | 0.58087 +/- 0.00568 |
| 0,002 | 0.26747 +/- 0.01657 |
| 0,001 | 0.16703 +/- 0.00309 |
| 0.0005 | 0.09803 +/- 0.000802 |

**Figure S13. A**. Linear fit of ATTO-488 carboxy absorbance values using solutions of 10$^{-2}$ μmol/mL, 8 10$^{-3}$ μmol/mL, 6 10$^{-3}$ μmol/mL, 4 10$^{-3}$ μmol/mL, 2 10$^{-3}$ μmol/mL, 10$^{-3}$ μmol/mL and 5 10$^{-4}$ μmol/mL in PBS pH 7.6. **B**. Absorbance values of the ATTO-488 carboxy at 500 nm based on three technical replicates. Error bars correspond to the standard deviation of the mean.

#### Calculation of membrane (ATTO-655) and cargo (ATTO-488 carboxy) signal in bulk over time

Non-modified and 5% poly(DPA) modified liposome populations labelled with ATTO-644 and loaded with ATTO-488 carboxy (10$^{-2}$ μmol/mL, $C_0$) in 10 mM PBS pH 7.6 were prepared. The absorbance of ATTO-488 carboxy at 500 nm and the fluorescence of ATTO-655 at 647 nm were measured immediately after preparation and after 5, 10 and 15 days of storage at 4 °c. The absorbance and fluorescence measurements were conducted in triplicates, and the absorbance was converted to ATTO-488 carboxy concentration using the formula of linear regression. The liposome populations were purified *via* dialysis in 10 mM PBS pH 7.6 before every measurement in order for the leaked ATTO-488 carboxy to be removed.

**Table S1. ATTO-488 carboxy leakage calculations on plate reader assay**

| Liposome | Day | Abs at 500 nm | ATTO-488 concentration (μmol/mL) | Leakage (%) |
|---|---|---|---|---|
| Non-modified | 0 | 0.5601 +/- 0.0036 | 4.02 10$^{-3}$ +/- 2.52 10$^{-4}$ | - |
| Non-modified | 5 | 0.3849 +/- 0.0192 | 2.69 10$^{-3}$ +/- 1.45 10$^{-4}$ | 33.1 |
| Non-modified | 10 | 0.2820 +/- 0.0041 | 1.90 10$^{-3}$ +/- 0.33 10$^{-4}$ | 52.7 |
| Non-modified | 15 | 0.2112 +/- 0.0089 | 1.36 10$^{-3}$ +/- 0.70 10$^{-4}$ | 66.2 |
| 5% poly(DPA) modified | 0 | 0.8385 +/- 0.0177 | 6.15 10$^{-3}$ +/- 1.3 10$^{-4}$ | - |
| 5% poly(DPA) modified | 5 | 0.7360 +/- 0.0103 | 5.37 10$^{-3}$ +/- 0.76 10$^{-4}$ | 12.7 |
| 5% poly(DPA) modified | 10 | 0.7016 +/- 0.0024 | 5.10 10$^{-3}$ +/- 0.21 10$^{-4}$ | 17.1 |
| 5% poly(DPA) modified | 15 | 0.6849 +/- 0.0120 | 4.98 10$^{-3}$ +/- 0.91 10$^{-4}$ | 19.1 |

**Table S2. ATTO-655 signal on plate reader assay**

| Liposome | Day | Fluorescence at 647 nm | Normalized signal to day 0 |
|---|---|---|---|
| Non-modified | 0 | 1959508.6 +/- 30919.9 | 1 +/- 0.014 |
| Non-modified | 5 | 1322584.6 +/- 16444.4 | 0.676 +/- 0.009 |
| Non-modified | 10 | 1363182.6 +/- 262732.8 | 0.693 +/- 0.131 |
| Non-modified | 15 | 1019874.0 +/- 41273.2 | 0.517 +/- 0.017 |
| 5% poly(DPA) modified | 0 | 4889571.3 +/- 68019.3 | 1 +/- 0.013 |
| 5% poly(DPA) modified | 5 | 5049702.3+/- 43409.4 | 1.033 +/- 0.008 |
| 5% poly(DPA) modified | 10 | 4854462.0 +/- 16541.5 | 0.992 +/- 0.003 |
| 5% poly(DPA) modified | 15 | 4650156.0 +/- 232709.4 | 0.951 +/- 0.047 |

**Calculation of ATTO-488 carboxy leakage using a single particle assay**

Non-modified and 5% poly(DPA) modified liposome populations loaded with ATTO-488 carboxy were imaged on TIRF immediately after preparation and 5 days after storage at 4 °C. The intensity loss in the blue channel after 5 days of storage was evaluated. The liposome samples were purified *via* dialysis in 10 mM PBS pH 7.6 before every measurement in order for the leaked ATTO-488 carboxy to be removed.

**Table S3. ATTO-488 carboxy intensity loss in the single particle assay**

| Liposome | Day | Mean Intensity in the blue channel | Leakage (%) |
|---|---|---|---|
| Non-modified | 1 | 6719.0 | - |
| Non-modified | 5 | 4211.0 | 33 |
| 5% poly(DPA) modified | 1 | 4948.0 | - |
| 5% poly(DPA) modified | 5 | 4243.0 | 14 |

**8. Liposome populations size**

The hydrodynamic diameter of the liposome populations was measured on DLS in 10 mM PBS pH 7.6 (C=2.5 μg/mL).

**Table S4**. Average size of liposome populations and PDI values based on four technical replicates, where error bars correspond to the standard deviation of all the detected particles.

| Liposome modification | Cargo | Mean Diameter (nm) | PDI |
|---|---|---|---|
| Non-modified | ATTO-488 carboxy | 147.1 +/- 64.57 | 0.14 |
| 5% poly(DPA) | ATTO-488 carboxy | 156.5 +/- 58.85 | 0.19 |
| 5% poly(DPA) | - | 111.9 +/- 54.44 | 0.24 |
| 1% poly(DPA) | ATTO-488 carboxy | 164.4 +/- 46.35 | 0.08 |
| 1% poly(DPA) | - | 174.9 +/- 51.5 | 0.09 |
| 1% poly(DEAEMA) | ATTO-488 carboxy | 183.9 +/- 61.99 | 0.11 |
| 1% poly(DEAEMA) | - | 169.1 +/- 51.6 | 0.09 |
| 5% poly(DPA) | Methylene blue | 175.8 +/- 63.23 | 0.12 |
| 5% poly(DPA) | siRNA | 155.0 +/- 62.59 | 0.16 |
| 5% poly(DPA) | BSA | 134.1 +/- 92.77 | 0.47 |

**Effect of the pH on the hydrodynamic diameter of the liposome populations**

The hydrodynamic diameter of all the prepared liposome populations was measured on DLS in 10 mM PBS pH 7.6 (C=2.5 μg/mL) and in 156 mM sodium acetate buffer pH 5.2 (C=2.5 μg/mL). The pH drop of the liposome populations (prepared in PBS pH 7.6) was achieved by dialysis in 156 mM sodium acetate buffer pH 5.2, using dialysis bags (Spectra/Por® Biotech Regenerated Cellulose Dialysis Membranes, MWCO 8-10 kDa).

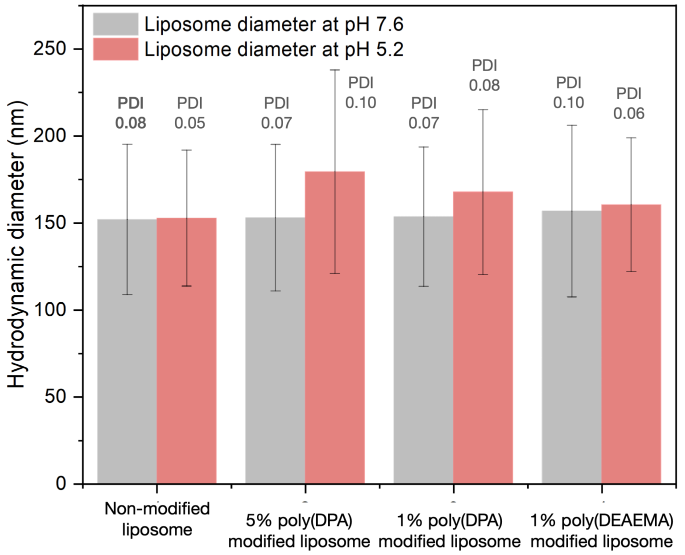


**Figure S14**. Average size of liposome populations and PDI values at pH 7.6 and pH 5.2 based on four technical replicates, where error bars correspond to the standard deviation of all the detected particles.

### 9. Single cell assay for d2eGFP knockdown

HEK293 cells stably expressing d2eGFP were established by transducing cells with a lentiviral vector encoding d2eGFP. The lentiviral particles were produced by transfecting the HEK293T packaging cell line with pMD2.G, psPAX2, and pLV-CMV-d2eGFP plasmid using Lipofectamine 3000 according to the manufacturer's protocol. HEK293 cells stably expressing d2eGFP were transferred to a preheated microscopy incubation chamber, and 5 positions with evenly distributed cells were selected for each experimental condition (cells in the absence of particles, and cells treated with siRNA loaded 5% poly(DPA) modified liposomes). Immediately before starting image acquisition, of 1 μL of liposome solution of a concentration of 1 mg/mL loaded with siRNA targeting d2eGFP were added dropwise to the medium of the wells of interest. Seven z-plane images with 3 μm z-spacing were acquired per position at 2.5 min intervals. Images were acquired for 10 h in total. For the excitation of the d2eGFP, 488nm laser was used with a 50,04 ms exposure time. Cells that didn't undergo cell division were selected for our analysis. Cell segmentation was performed using Cellpose software and tracking using a foundational mode (SAM2) which from a click on the selected cell at the first frame, SAM2 segmented the cell and tracked it for all further frames. Three out of six selected tracked cells in the control experiments showed increasing eGFP signal over time, while upon treatment with 5% ply(DPA) modified liposomes loaded with eGFP-targeting siRNA, six out of eight selected tracked cells showed a clear eGFP signal decrease.

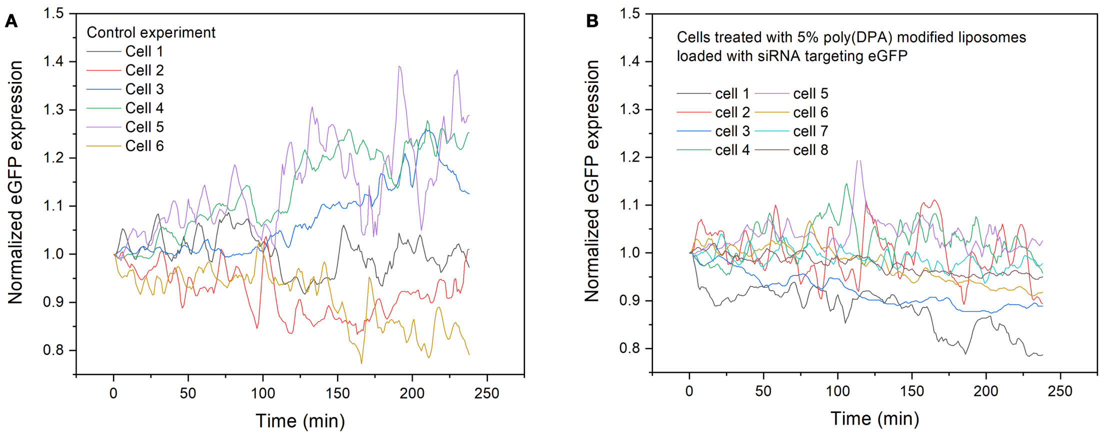


**Figure S15**. Selected traces normalised for each of the above experimental condition. Each time point intensity represents the median intensity of the cell mask normalized to the mean value of the 3 first points of a trace. Intensity traces were denoised with a uniform convolution window of length 5. **A**. Control experiment in the absence of particles. **B**. Cells were treated with 5% poly(DPA) modified liposomes loaded with siRNA targeting d2eGFP.
